# Long-term non-trophic effects of large herbivores on plant diversity are underestimated

**DOI:** 10.1101/2024.08.13.607836

**Authors:** Qingqing Chen, Jan P. Bakker, Juan Alberti, Elisabeth S. Bakker, Christian Smit, Han Olff

## Abstract

The positive effects of large herbivores on plant diversity in grasslands have so far been mainly attributed to increased light availability or suppressed dominance, and thus to the consequences of aboveground biomass consumption (trophic effects). However, these insights are mainly derived from short-term experiments. Using a 46-year experiment in a salt marsh, comparing cattle grazing, mowing (as a proxy of aboveground consumption) and the ungrazed control, we found that the non-trophic effects (e.g. trampling, deposition of urine and dung) of large herbivores on plant diversity increased over time, exceeding that of the trophic effects after 23 years. This long-term accumulation of non-trophic effects through slow ecosystem-level feedback highlights the sustainability of using low to moderate densities of large herbivores to conserve plant diversity. Our results emphasize the need for the conservation and re-introduction of large herbivores, domestic or wild, to sustain long-term grassland plant diversity.

## Introduction

Large herbivores promote plant diversity in grasslands worldwide (Olff & Ritchie 1998; Bakker *et al*. 2006; Borer *et al*. 2014; Davidson *et al*. 2017). Past studies conclude that herbivores mostly do this by increasing light availability at the ground level (Borer *et al*. 2014), or by suppressing dominant plant species (Mortensen *et al*. 2017; Koerner *et al*. 2018). These changes are often attributed to the aboveground biomass consumption (i.e. trophic effects) (e.g. Kohler et al. 2004; Mikola et al. 2009). However, these conclusions are mainly derived from short-term experiments, and it is so far unclear whether these prime effects of large herbivores – promoting plant diversity via trophic effects – also hold in the long term.

In addition to the trophic effects, large herbivores can also affect plant diversity via non-trophic effects, for instance, trampling, deposition of urine and dung (Kobayashi et al. 1997; Bakker & Olff 2003; Ludvíková et al. 2014; Van Klink *et al*. 2015; Lezama & Paruelo 2016). Trampling creates frequent small disturbances, which increases germination gaps for new species (Bullock et al. 1995; Ludvíková et al. 2014). Trampling also increases habitat heterogeneity (e.g. er content, microclimate). Similarly, local deposition of urine and dung increases heterogeneity in nutrient availability (Dai 2000; Mikola et al. 2009; Gillet et al. 2010). A heterogeneous environment allows more plant species to coexist than a homogeneous one (Vivian-smith 1997; Lundholm & Larson 2003; Davies et al. 2005). In addition, these non-trophic effects tend to accumulate over time (Mikola et al. 2009; Elschot et al. 2015; Van Klink et al. 2015), suggesting that short-term studies may underestimate their importance. However, really long-term evaluations (time scale of several decades) of the relative importance of the trophic versus non-trophic effects of herbivores on plant diversity have until now not been available. Here, we use a 46-year experiment in which we explore the long-term changes in the relative importance of the trophic (removal of biomass) and non-trophic (trampling + deposition of urine and dung) effects of large herbivores on plant diversity.

## Materials and Methods

### Study system

The experiment was conducted in the natural salt marsh of the barrier island of Schiermonnikoog (53°30’ N, 6°10’ E), the Netherlands (Bakker 1989). The average annual temperature is 9 °C, and average annual rainfall is 807 mm (data from www.knmi.nl). In this ecosystem, a natural succession gradient is present; the western part of the salt marsh has undergone more than 100 years’ succession (Olff *et al*. 1997), and is dominated by the tall late successional grass, *Elytrigia atherica*, when cattle grazing is absent. Primary productivity (measured inside herbivore exclosure) is high (1119.8 ± 201.4 g dw m^-2^ year^-1^; mean ± 1 se; measured in 2018) in this area, probably explaining why the dominance of *E. atherica* leads to a decline in plant diversity, a phenomenon widely observed in salt marshes across Europe (Pétillon *et al*. 2005; Milotić *et al*. 2010; Veeneklaas *et al*. 2013; Wanner *et al*. 2014; Rupprecht *et al*. 2015).

### Experimental design

The part of the salt marsh where the study was performed, ca. 32 ha, has been subject to cattle grazing up to 1958. Grazing stopped in 1958, which led to the dominance of *E. atherica* and decreased plant diversity over the following 10 years (Bakker 1989). In order to reverse this trend, cattle grazing with heifers restarted in 1972. Cattle graze from May to November in this area, after which they are taken out by the farmers and moved indoors during the storm season. Stocking density reduced from 1.5 to 0.5 heads ha^-1^from 1993 onwards, as the potential area that could be grazed expanded (Bakker *et al*. 1993) (Fig. S1). Cattle graze the vegetation to ca. 2 cm (Q. Chen personal observation). However, in this area, grazing also promotes heterogeneity in vegetation structure – patches of short grazing-tolerant plants alternate with those of tall grazing-avoiding plant species (Howison *et al*. 2017). The second author (JPB) established four exclosures to monitor the effects of cattle grazing on the plant communities in 1972. Exclosures (ca. 8 m × 42 m) were constructed with two electrical strands running 0.5 and 1 m above the ground. Exclosures were part of the design of four blocks, separated by at least 100 m. Each block contained: 1) cattle grazing, 2) late season mowing and 3) herbivore exclusion, that is an ungrazed control (detailed experimental design shown in Fig. S1). Mowing treatments were established inside each exclosure with an area of ca. 18 m^2^. The vegetation was mown to 2 cm height using a brush cutter, and the cut vegetation was raked, weighed and removed. The plots were mown annually in late August or early September (late season mowing) at peak standing crop (Bakker & De Vries 1992). Previous studies have used repeated mowing to simulate the trophic effects of large herbivore (Kohler et al. 2004; Mikola et al. 2009; Van Klink et al. 2015; Lezama & Paruelo 2016). Following that, we used the data from the late season mowing as a proxy of trophic effects of cattle grazing. We compared late season mowing with the early season mowing (usually in June) and the both early and late season mowing (Fig. S2A). We found that, averaged over the 28 occasions of recordings during the 46-year experiment, late season mowing removed the biomass the most (early season mowing: 215.9 ± 13.53 g dw m^-2^ mean ± 1 se; late season mowing: 281.5 ± 15.46; both early and late season mowing: 231.8 ± 15.25). More importantly, the amount of aboveground biomass removed by late season mowing was similar to the amount removed by cattle grazing (Fig. S2B).

We marked one permanent plot (2 m × 2 m) in each treatment and block in 1972. We recorded plant species occurrence and abundance in the permanent plots before the late season mowing from 1972 to 2017, 33 occasions of recordings in total (annually from 1972 to 1980 and 1984 to 2001, and 2004, 2006, 2009, 2013, 2017). Plant diversity was measured as number of species within the permanent plots. The abundance (percent cover) was estimated using the decimal scale of Londo (1976), then transformed to percent cover using a standard procedure (Chen *et al*. 2021). The majority of the recordings was done by a skilled field assistant, Y. de Vries. A total of 48 plant species were recorded during the 46-year experiment (a full species list is given in Table S1).

Before the late season mowing, we measured proportion of photosynthetically active radiation (PAR, μmole photons m^-2^ s^-1^) on a sunny day (September 2018, between 12:00 and 14:00, approximately solar noon) using a light sensor (Skye, UK). We took four measurements per permanent plot. For each measurement, we simultaneously measured PAR at ground level (ca. 3.8 cm) and above vegetation (ca. 50 -100 cm). We calculated light availability as the PAR reaching ground level compared to that of the above vegetation. Four measurements were averaged per permanent plot.

## Data analysis

### Plant diversity and species turnover

We analyzed change in plant diversity (i.e. number of species) in different treatments over time. Grazing affects plant diversity via changing species gain (colonization) and species loss (extinction) (Olff & Ritchie 1998). Therefore, we partitioned plant diversity into species gain and loss. Cumulative species gain or loss was calculated as: number of species gained or species lost divided by the total number of species recorded in a given year compared with the starting year 1972 using the R package codyn (Hallett *et al*. 2016). To explore whether treatments promoted particular functional groups of plant species (number and abundance), we classified them into forbs, graminoids, legumes and woody species according to their life forms (see Table S1 for classification of life form for each species).

### Dominance

We explored the underlying mechanism of changes in plant diversity via reducing dominance over time (Koerner *et al*. 2018). Dominance was measured as the Berger-Parker dominance index, i.e. the proportional abundance of the most abundant species. In addition, we classified plant species into rare, frequent, common and abundant species according to their abundances. Rare: percent cover in any permanent plot in any year ≤ 1; frequent: > 1 and ≤ 20; common: > 20 and ≤ 50; abundant: > 50. We explored whether treatments promoted particular functional groups in terms of their abundances.

### Halophytes

Given that herbivore trampling induces anoxic conditions, which could favor halophyte species (Van Klink et al. 2015), we also analyzed changes of this functional group. We analyzed changes in the number and abundance of halophyte species over time. Halophytes are plant species well adapted to anoxic condition (Bakker et al. 2002; Van Klink et al. 2015), which is usually induced by inundation in salt marshes, but also by soil compaction due to trampling (Schrama et al. 2013a). We classified halophytes according to Bakker et al. (2002). See Table S1 for a list of halophyte species.

We calculated the total effects of large herbivores on plant community properties such as plant diversity as a property in the grazing treatment minus that in the ungrazed control, the trophic effects as the mowing treatment minus the ungrazed control, and the non-trophic effects as the grazing treatment minus the mowing treatment. We evaluated the total effects of large herbivores in one model, and the trophic and non-trophic effects in another model, using generalized additive mixed models (gamm) from the package mgcv (Wood 2017). In all models, permanent plot was a random variable, temporal autocorrelation was adjusted using the corCAR1 model, due to the unevenly-spaced sampling. Temperature, rainfall and sea level change were not correlated with change in plant diversity over time, therefore were not included in the models (Fig. S3; online supporting text). Legumes and woody species were very rare, therefore, we only analyzed the probability of their presence in different treatments. We used the binomial distribution in these two models. Autocorrelation was not included in the model of the abundance of legumes, as the AIC increased when including the autocorrelation. We extracted the parametric coefficients and smooth terms from these models to assess the overall effects (average across 46 years) and over-time effects. Data were analyzed in R 3.5.3 (R Core Team, 2019).

## Results

### Plant diversity and species turnover

Averaged over all years, large herbivores increased plant diversity by 5.91 species per 4 m^2^ compared with the ungrazed control (estimated from gamm models; Table S2). Non-trophic and trophic effects contributed 2.81 and 2.99 plant species to this increase, respectively (Table S2). Large herbivores promoted plant diversity over time (F = 3.21, *p* = 0.0200; Table S3), particularly in the first half (23 years) of the experiment (Fig. 1A). The contribution of non-trophic effects on plant diversity increased over time (F = 2.52, p = 0.0547; Table S3), and exceeded the trophic effects 23 years after the start of the experiment (Fig.1B). Averaged over all years, large herbivores promoted plant diversity via increasing species gain and decreasing species loss (Fig. 1C, E; Table S2). Large herbivores decreased species loss over time (F = 7, p = 0.0011; Table S3), and the contribution of the non-trophic effects increased and exceeded that of the trophic effects 27 years after the start of the experiment (F = 10.82, p = 0.0011; Fig. 1F; Table S3). Whereas the trophic effects decreased cumulative species gain over time (F = 4.04, p = 0.04; Fig. 1D; Table S3), and their contribution to the total effects of large herbivores were exceeded by the non-trophic effects 13 years after the start of the experiment (Fig. 1D). Large herbivores promoted the number and abundance of forbs over time, and the contribution of non-trophic effects increased over time (Fig. S4B; S5B; Table S3; online supporting text). Large herbivores promoted the number of species but decreased the abundance of graminoids over time. The contribution of the non-trophic effects increased in promoting the number of species of graminoids. However, the trophic effects contributed more than the non-trophic effects to the total effects of large herbivores in decreasing the abundance of graminoids after 27 years (Fig. S4D; S5D; Table S3; online supporting text).

**Fig. 1.**
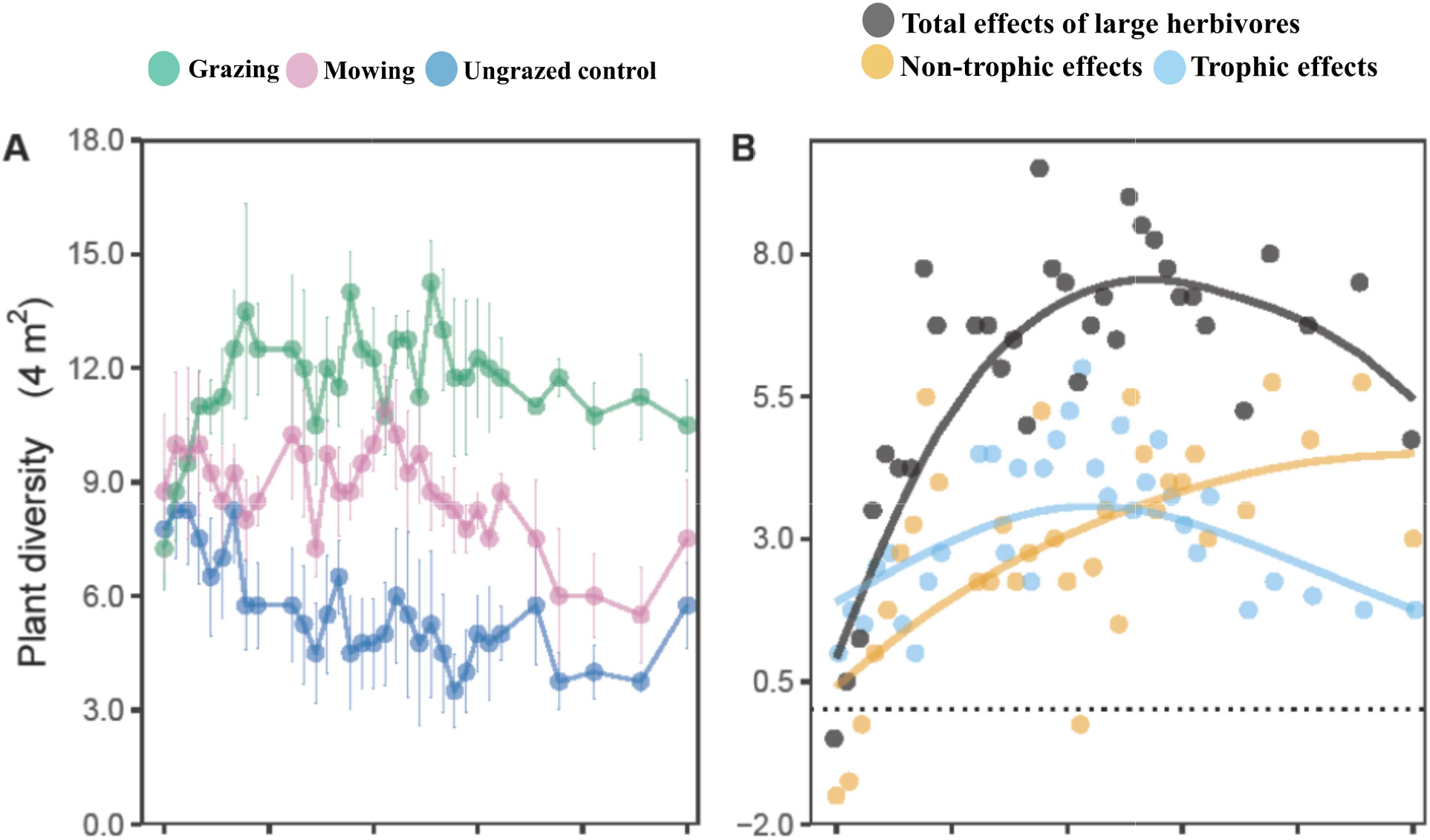
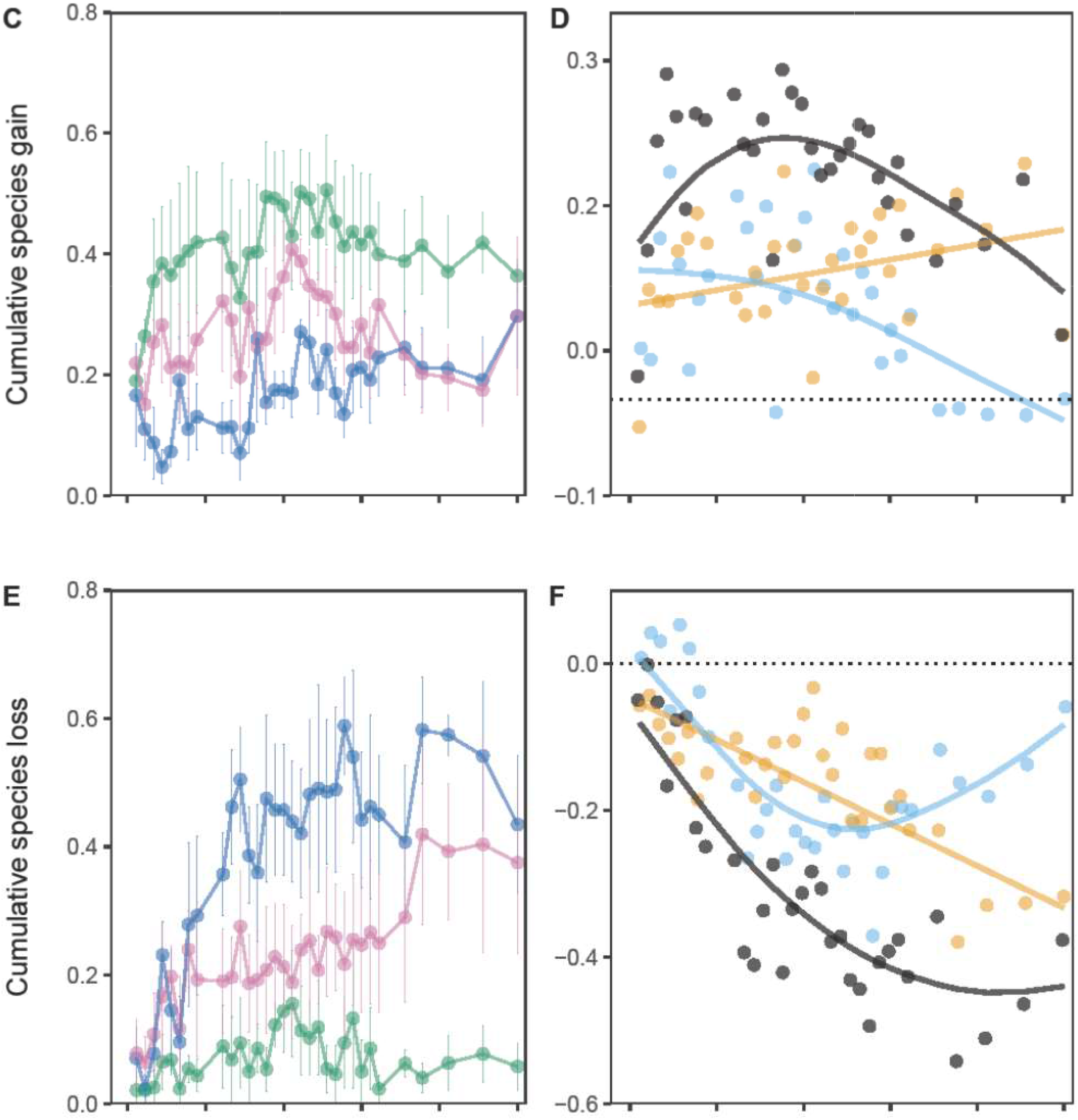
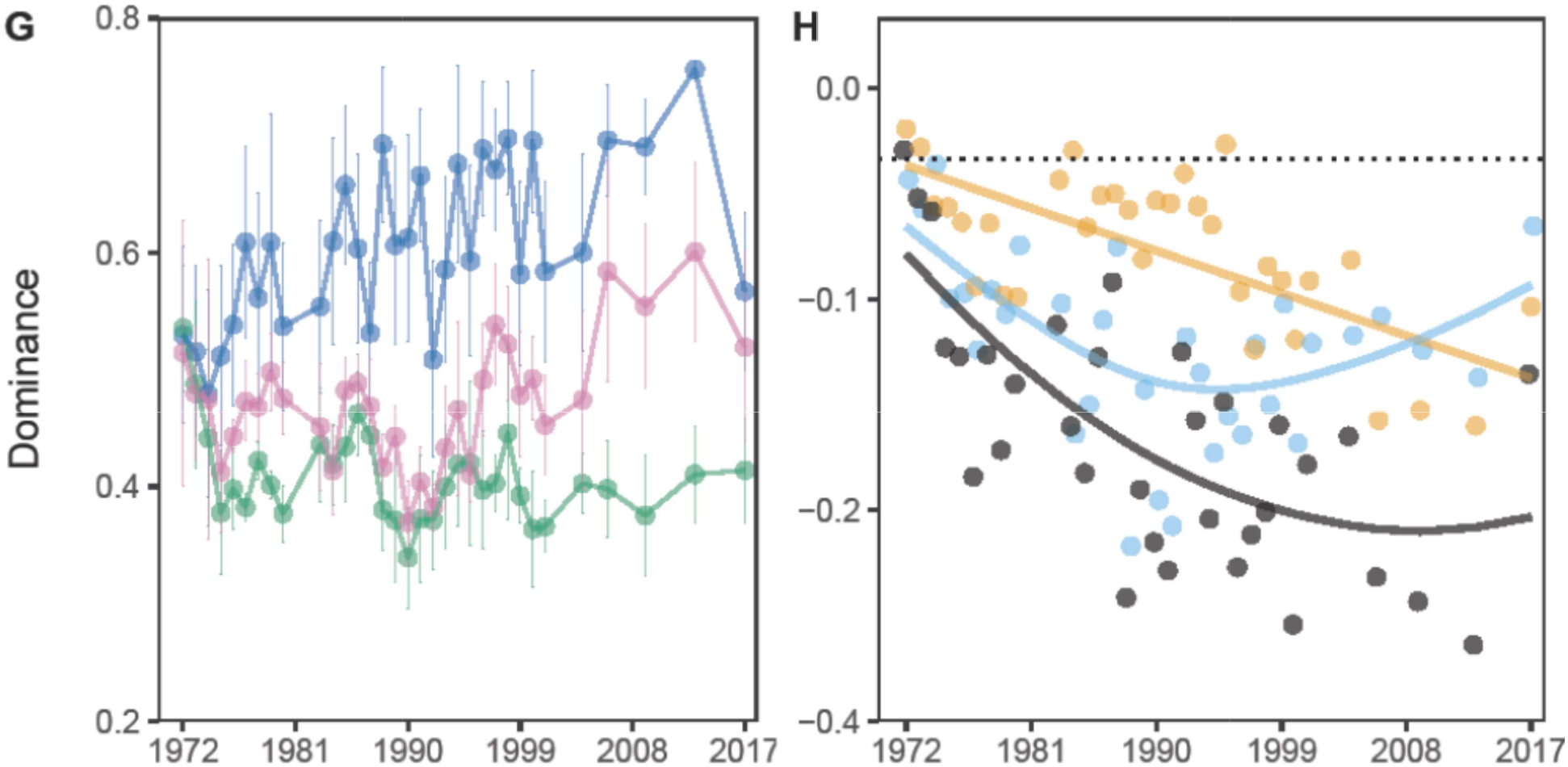
Plant diversity, species gain, species loss and dominance during the 46-year experiment. Large herbivores promoted plant diversity over time, and the contribution of non-trophic effects increased and exceeded that of the trophic effects 23 years after the start of the experiment (A, B). Large herbivores decreased species loss over time, and the contribution of non-trophic effects exceeded that of the trophic effects on species gain and loss 13 and 27 years after the start of the experiment, respectively (C, D, E, F). Large herbivores decreased dominance over time, and the contribution of non-trophic effects increased (G, H). Cumulative species gain or loss was calculated as the number of species gained or lost divided by the total number of species recorded in a given year compared with the starting year 1972. Dominance was expressed as the Berger-Parker dominance index, i.e. the proportional abundance of the most abundant species in a given plot. Total effects of large herbivores: the grazing treatment minus the ungrazed control; non-trophic effects: the grazing treatment minus the mowing treatment; trophic effects: the mowing treatment minus the ungrazed control. Dots show means of four blocks. Bars reflect ±1se. Lines in B, D, F, H are fitted with generalized additive mixed models (Table S3).

### Dominance

Averaged over all years, large herbivores decreased plant species dominance by 20 % (Table S2). Non-trophic and trophic effects contributed 7 % and 13 % to this decrease, respectively (Table S2). Large herbivores decreased dominance over time (*F* = 7.39, *p* = 0.0007; Table S3), and the contribution of non-trophic effects increased (*F* = 6.1, *p* = 0.0141; Fig. 1H; Table S3). Large herbivores decreased dominance by promoting the number of rare, frequent, and common, but not dominant species over time, and the contribution of non-trophic effects increased in the frequent and common species (Fig. S6; Table S3). By contrast, dominance increased in the ungrazed control, which was mainly due to the expansion of the tall late successional grass *E. atherica* (Fig. S7A, B; online supporting text). Trophic effects alone (i.e. late season mowing) led to dominance of another grass, *Festuca rubra* (Fig. S7C, D), forming a dense sward, which may explain the ultimate decline in plant diversity in this treatment (Fig.1A,1G, S8).

### Light availability

Forty-six years after the start of the experiment, the percentage of photosynthetically active radiation reaching ground level in the grazed treatment increased by more than 10-fold compared with the ungrazed control (ungrazed control: 7 ± 1 %; grazing: 73 ± 22 %; mowing: 33 ± 20 %; mean ± 1se). Non-trophic effects contributed ca. 61 % to the total effects of large herbivores on light availability.

### Halophytes

Averaged over all years, large herbivores increased 2.93 halophytes, and non-trophic effects contributed 2.41 to this increase. Also, large herbivores increased the abundance of halophytes by 14 %, which was entirely attributed to their non-trophic effects (Table S2). In addition, large herbivores promoted the number and abundance of halophytes over time, which was mostly attributed to their non-trophic effects (Fig. 2; Table S3).

**Fig. 2.**
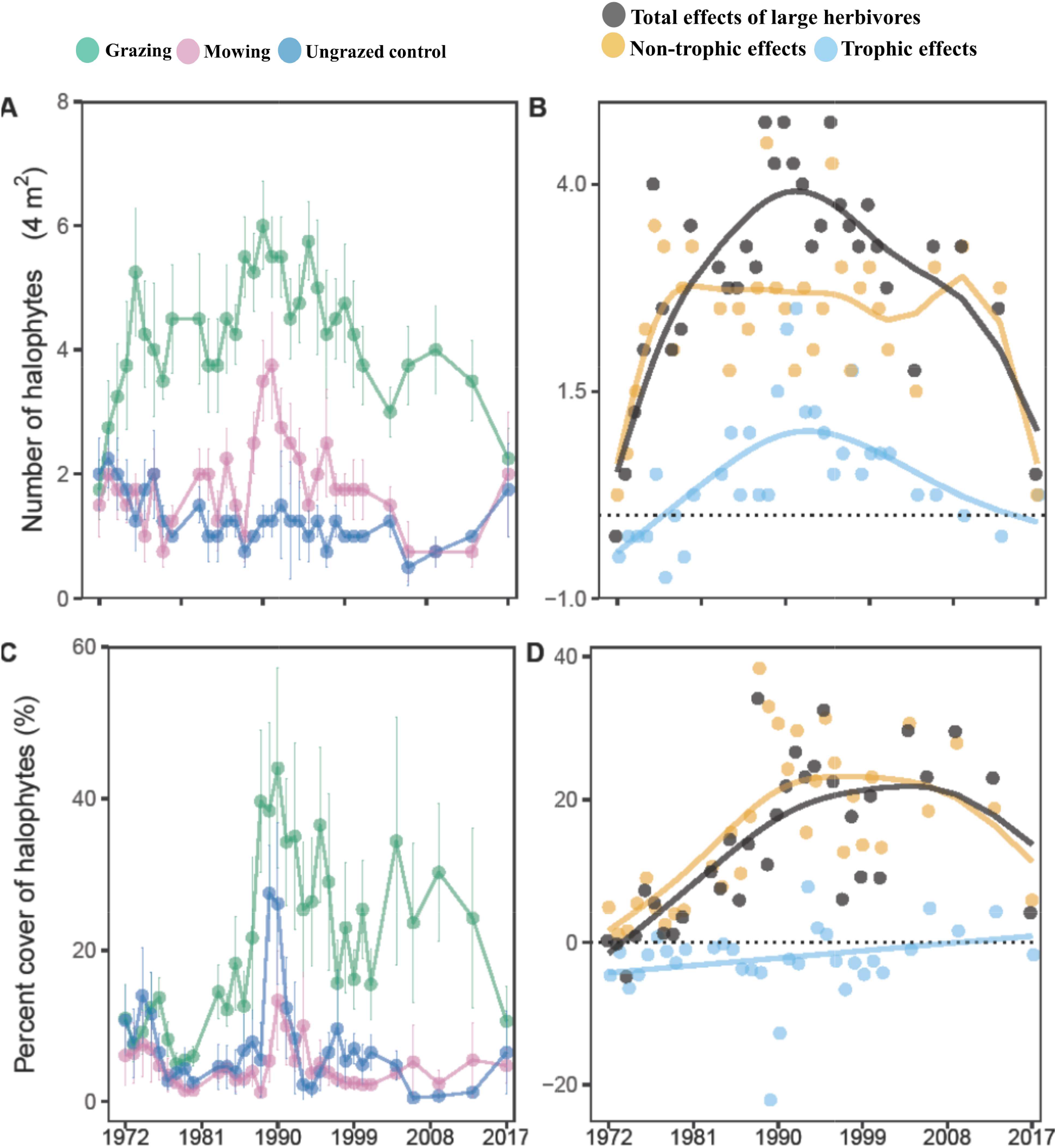
Number and abundance of halophytes. Large herbivores promoted the number and abundance of halophytes over time, which could mostly be attributed to the non-trophic effects. Total effects of large herbivores: the grazing treatment minus the ungrazed control; non-trophic effects: the grazing treatment minus the mowing treatment; trophic effects: the mowing treatment minus the ungrazed control. Dots show means of four blocks. Bars reflect ±1se. Lines in B were fitted with generalized additive mixed models (Table S3).

## Discussion

Our unique 46-year experiment suggests that in the first half (23 years) of the experiment the effects of large herbivores on plant diversity were more attributed to their trophic effects (biomass removal), simulated with late season mowing. However, the non-trophic effects increased in importance over time, and outweighed the trophic effects in the second half of the experiment, despite the reduction in stocking density from 1.5 to 0.5 heads ha^-1^. Large herbivores increased plant diversity via reducing species loss over time.

The non-trophic effects played an important, but so far overlooked, role in promoting plant diversity, particularly in the long term. Our results provide a good explanation for the low importance of the non-trophic effects on plant communities in short-term grazing experiments (1 year in Kohler et al. 2004; 3 years in Mikola et al. 2009). Consistent with previous analyses, large herbivores likely promote plant diversity via decreasing dominance (Koerner *et al*. 2018), and increasing light availability (Borer *et al*. 2014). For instance, we found that large herbivores increased photosynthetically active radiation reaching ground level by more than 10-fold and similar to the effects observed 17 years after the start of this experiment (Bakker & De Vries 1992). Large herbivores did so likely through reducing abundance of dominant plants and overall aboveground biomass but not through reducing plant height (Chen *et al*. 2021). However, the trophic effects decreased, whereas non-trophic effects increased and exceeding that of trophic effects on plant diversity in the second half of the experiment. Large herbivores may create open gaps with bare soil, which could serve as a regeneration niche to promote plant colonization and diversity (Bakker 1985; Bakker *et al*. 1985). Overall, large herbivores did not impact percent cover of bareground. But in the first half of the experiment, large herbivores slightly increased percent cover of bareground, which was primarily attributed to non-trophic effects (Fig. S9).

This increased open gaps with bare soil under herbivore grazing may promote plant colonization and species gain at the first 15 years of the experiment.

The non-trophic effects of large herbivores can alter belowground variables, which likely feedback to the aboveground variables. Past studies in this system found that long-term grazing alters soil properties such as increasing soil bulk density, reducing soil redox potential, and soil macrofauna abundances (Schrama *et al*. 2013a). This alteration of soil properties via trampling, and to a lesser extent, local deposition of urine and dung, subsequently alters nutrient cycling (e.g. decreasing nitrogen mineralization and increasing carbon stock) (Schrama *et al*. 2013a; Elschot *et al*. 2015). In addition, this alteration of soil properties impacts plant communities, for instance, promoting the number and abundance of halophytes. Halophytes are good indicators for anoxic conditions induced by trampling (Bakker *et al*. 2002; Van Klink *et al*. 2015). Although anoxic conditions may be confounded with inundation in salt marshes. In this experiment, the increased number and abundance of halophytes were mainly induced by trampling as halophytes did not increase in the mowing or the ungrazed control treatment. Previous studies in this area also showed that inundation frequency and salinity play an insignificant role in plant-plant interactions and community structure compared with cattle grazing (Bakker 1985; Bakker *et al*. 1985; Howison *et al*. 2015). Further, this non-trophic effects tend to accumulate over time, as the elevation decreased in this area over time (measured from 1994 to 2017), primarily driven by trampling (Elschot *et al*. 2015). Therefore, we expect the increased importance of the non-trophic effects of large herbivores over time observed in this system to be also important in grasslands with fine soil texture, resulting in compactable soil (Schrama *et al*. 2013b, a).

The late season mowing can be a good proxy of the trophic effects of large herbivores in this system. We verified that the amount of aboveground biomass removed by mowing was similar to the amount consumed by cattle (Fig. S2B). Surprisingly, few studies verify this when using mowing as a proxy of trophic effects of herbivores (e.g. Kohler *et al*. 2004; Ludvíková *et al*.

2014; Van Klink *et al*. 2015; but see Mikola *et al*. 2009). It is not surprising that mowing can have a more positive effect on plant diversity than grazing, if mowing removes more aboveground biomass, thus increasing light availability or decreasing dominance to a greater extent. Mowing can, however, still be different from the trophic effects of herbivores. For instance, mowing removes aboveground biomass at one time, while herbivores consume vegetation throughout the whole grazing season. Mikola et al. (2009) compared cattle grazing (4 to 5 times rotation per year) to mowing (4 to 5 times at the beginning of each rotation), and found that removal of aboveground biomass (the trophic effects) was the major driver of changes in plant variables such as aboveground biomass, shoot N and P concentration. However, in this 3-year experiment, although soil inorganic N concentration increased over the years, soil moisture, and soil density was similar in the grazing, mowing and ungrazed control treatments, suggesting that the non-trophic effects has not yet taken place in this short-term experiment.Also, mowing imposes a uniform disturbance to all vegetation, while grazing imposes a more selective one (Tälle *et al*. 2016). Mowing, on the other hand, may promote graminoids more than forbs, possibly because the meristems of graminoids are situated at the base of tillers, while forbs have terminal meristems and may thus suffer more damage from grazing than graminoids. Indeed, we found that, in this graminoid-abundant system, grazing promoted the number and the abundance of forbs over time as compared with the mowing treatment (Fig. S4; S5). However, the abundance of woody species (*Artemisia maritima*) increased in the mowing treatment after 1999 (Fig. S5G, H; Table S3), suggesting that plants with the terminal meristems were not particularly suppressed by mowing in this system. In addition, the abundance of graminoids became similar in the grazing and mowing treatment, particularly in the end of the experiment (Fig. S5C).

Therefore, the late season mowing and cattle grazing treatment did not show distinct differences in vegetation selectivity. Taken together, the late season mowing can be a good proxy of the trophic effects of large herbivores in this system.

Our results have clear implications for conservation and management of plant diversity in salt marshes and grasslands. Using this long-term cattle grazing experiment as a model, we showed that low to moderate densities (usually relative to the productivity of the site) of large domestic herbivores, and probably also wild ones, would play a positive role in conserving plant diversity. However, patience is required from conservation managers, as the key results and underlying mechanisms take several decades to develop. Global livestock production is increasing (Thornton 2010; Fig. S10A). However, livestock (e.g. dairy cows) is increasingly being kept indoors (Mandel *et al*. 2016), fed by mown grasses and crops, particularly in the developed world. For instance, in Denmark, all livestock is kept indoors during the lifetime (Reijs *et al*. 2013). Grasslands are mowed or converted to croplands, and the non-trophic effects of large herbivores have thus been removed, which has contributed to the decline in plant diversity (O’Mara 2012; Tscharntke *et al*. 2012). Next to agricultural intensification, farmland abandonment has emerged in recent years (Fig. S10B), which has also caused a decline in plant diversity (Benayas *et al*. 2007). In Europe, the area of the abandoned farmland (agricultural grasslands) increased by more than 15 % since 1992 (Fig. S10B), and this is projected to increase in the next 30 years (Terres *et al*. 2015). Also, in natural grasslands, wild herbivores – particularly wild megaherbivores – have been declining dramatically worldwide (Ripple *et al*. 2015). Our study therefore emphasizes the benefit of reintroducing large herbivores (wild or domestic) in the abandoned grasslands with an evolutionary history of grazing. In addition, extensive pasture-based livestock farming should be encouraged and maintained if biodiversity is to be conserved in the long term.

## Supporting information

Supplementary file for long-term non-trophic effects

## Acknowledgments

We thank Natuurmonumenten for offering us the opportunity to work in the salt marsh of the island of Schiermonnikoog. We thank Y. de Vries for recording the vegetation over the years. We thank Ido Pen for his helpful suggestions for data analysis. We thank Yong-fei Bai for his constructive comments for our manuscript. QC is funded by CSC (China Council Scholarship). JA was supported by a Visitor’s Travel Grant (040.11.631) of the Netherlands Organisation for Scientific Research (NWO).

## Author contributions

JPB designed and conducted the experiments. CS and QC maintained the experiment and collected data since 2012, and 2016, respectively. QC, HO and JA conceived the idea, QC analyzed the data and wrote the manuscript. All authors contributed to revisions, and gave final approval for publication.

## Data and materials availability

data will be deposited in the Dryad Digital Repository once the manuscript gets accepted.

## References

Bakker, E.S. & Olff, H. (2003). Impact of different-sized herbivores on recruitment opportunities for subordinate herbs in grasslands. J. Veg. Sci., 14, 465–474.

Bakker, E.S., Ritchie, M.E., Olff, H., Milchunas, D.G. & Knops, J.M.H.H. (2006). Herbivore impact on grassland plant diversity depends on habitat productivity and herbivore size. Ecol. Lett., 9, 780–788.

Bakker, J.P. (1985). The impact of grazing on plant communities, plant populations and soil conditions on salt marshes. Vegetatio, 62, 391–398.

Bakker, J.P. (1989). Nature management by grazing and cutting: on the ecological significance of grazing and cutting regimes applied to restore species-rich grassland communities in the Netherlands. University of Groningen. Kluwer Academic Publishers, Dordrecht.

Bakker, J.P., Dijkstra, M. & Russchen, P.T. (1985). Dispersal, germination and early establishment of halophytes and glycophytes on a grazed and abandoned salt-marsh gradient. New Phytol., 101, 291–308.

Bakker, J.P., Esselink, P., Dijkema, K.S., Van Duin, W.E. & De Jong, D.J. (2002). Restoration of salt marshes in the Netherlands. Hydrobiologia, 478, 29–51.

Bakker, J.P., De Vlas, J. & Van Tooren, B.F. (1993). Uitbreiding begrazing van de Oosterkwelder op Schiermonnikoog. Levende Nat., 94, 118–122.

Bakker, J.P. & De Vries, Y. (1992). Germination and early establishment of lower salt-marsh species in grazed and mown salt marsh. J. Veg. Sci., 3, 247–252.

Benayas, J.M.R., Martins, A., Nicolau, J.M. & Schulz, J.J. (2007). Abandonment of agricultural land: An overview of drivers and consequences. CAB Rev. Perspect. Agric. Vet. Sci. Nutr. Nat. Resour., 2.

Borer, E.T., Seabloom, E.W., Gruner, D.S., Harpole, W.S., Hillebrand, H., Lind, E.M., et al. (2014). Herbivores and nutrients control grassland plant diversity via light limitation. Nature, 508, 517–520.

Bos, D., Bakker, J.P., De Vries, Y. & Van Lieshout, S. (2002). Long-term vegetation changes in experimentally grazed and ungrazed back-barrier marshes in the Wadden Sea. Appl. Veg. Sci., 5, 45–54.

Bullock, J.M., Hill, B.C., Silvertown, J., Sutton, M., Bullock, J.M., Hill, B.C., et al. (1995). Gap Colonization as a Source of Grassland Community Change : Effects of Gap Size and Grazing on the Rate and Mode of Colonization by Different Species. Oikos, 72, 273–282.

Chen, Q., Bakker, J. P., Alberti, J., Bakker, E. S., Smit, C., & Olff, H. (2021). Long-term cross-scale comparison of grazing and mowing on plant diversity and community composition in a salt-marsh system. Journal of Ecology, 109, 3737–3747. 10.1111/1365-2745.13753

Dai, X. (2000). Impact of Cattle Dung Deposition on the Distribution Pattern of Plant Species in an Alvar Limestone Grassland. J. Veg. Sci., 11, 715–724.

Davidson, K.E., Fowler, M.S., Skov, M.W., Doerr, S.H., Beaumont, N. & Griffin, J.N. (2017). Livestock grazing alters multiple ecosystem properties and services in salt marshes: a meta-analysis. J. Appl. Ecol., 54, 1395–1405.

Davies, K.F., Chesson, P., Harrison, S., Inouye, B.D., Melbourne, B.A. & Rice, K.J. (2005). Spatial Heterogeneity Explains the Scale Dependence of the Native-Exotic Diversity Relationship. Ecology, 86, 1602–1610.

Elschot, K., Bakker, J.P., Temmerman, S., Van De Koppel, J. & Bouma, T.J. (2015). Ecosystem engineering by large grazers enhances carbon stocks in a tidal salt marsh. Mar. Ecol. Prog. Ser., 537, 9–21.

Gillet, F., Kohler, F., Vandenberghe, C. & Buttler, A. (2010). Effect of dung deposition on small-scale patch structure and seasonal vegetation dynamics in mountain pastures. Agric. Ecosyst. Environ., 135, 34–41.

Hallett, L.M., Jones, S.K., MacDonald, A.A.M., Jones, M.B., Flynn, D.F.B., Ripplinger, J., et al. (2016). codyn: An r package of community dynamics metrics. Methods Ecol. Evol., 7, 1146–1151.

Howison, R.A., Olff, H., Van De Koppel, J. & Smit, C. (2017). Biotically driven vegetation mosaics in grazing ecosystems: the battle between bioturbation and biocompaction. Ecol. Monogr., 87, 363–378.

Howison, R.A., Olff, H., Steever, R. & Smit, C. (2015). Large herbivores change the direction of interactions within plant communities along a salt marsh stress gradient. J. Veg. Sci., 26, 1159–1170.

Van Klink, R., Schrama, M., Nolte, S., Bakker, J.P., WallisDeVries, M.F. & Berg, M.P. (2015). Defoliation and Soil Compaction Jointly Drive Large-Herbivore Grazing Effects on Plants and Soil Arthropods on Clay Soil. Ecosystems, 18, 671–685.

Kobayashi, T., Hori, Y. & Nomoto, N. (1997). Effects of trampling and vegetation removal on species diversity and micro-environment under different shade conditions. J. Veg. Sci., 8, 873–880.

Koerner, S.E., Smith, M.D., Burkepile, D.E., Hanan, N.P., Avolio, M.L., Collins, S.L., et al. (2018). Change in dominance determines herbivore effects on plant biodiversity. Nat. Ecol. Evol., 2, 1925–1932.

Kohler, F., Gillet, F., Gobat, J.M. & Buttler, A. (2004). Seasonal vegetation changes in mountain pastures due to simulated effects of cattle grazing. J. Veg. Sci., 15, 143–150.

Lezama, F. & Paruelo, J.M. (2016). Disentangling grazing effects: trampling, defoliation and urine deposition. Appl. Veg. Sci., 19, 557–566.

Londo, G. (1976). The decimal scale for releves of permanent quadrats. Vegetatio, 33(1), 61–64. 10.1007/BF00055300

Ludvíková, V., Pavlů, V. V., Gaisler, J., Hejcman, M. & Pavlů, L. (2014). Long term defoliation by cattle grazing with and without trampling differently affects soil penetration resistance and plant species composition in Agrostis capillaris grassland. Agric. Ecosyst. Environ., 197, 204–211.

Lundholm, J.T. & Larson, D.W. (2003). Relationships between Spatial Environmental Heterogeneity and Plant Species Diversity on a Limestone Pavement. Ecography (Cop.)., 26, 715–722.

Mandel, R., Whay, H.R., Klement, E. & Nicol, C.J. (2016). Invited review: Environmental enrichment of dairy cows and calves in indoor housing. J. Dairy Sci., 99, 1695–1715.

Mikola, J., Setälä, H., Virkajärvi, P., Saarijärvi, K., Ilmarinen, K., Voigt, W., et al. (2009). Defoliation and patchy nutrient return drive grazing effects on plant and soil properties in a dairy cow pasture. Ecol. Monogr., 79, 221–244.

Milotić, T., Erfanzadeh, R., Pétillon, J., Maelfait, J.P. & Hoffmann, M. (2010). Short-term impact of grazing by sheep on vegetation dynamics in a newly created salt-marsh site. Grass Forage Sci., 65, 121–132.

Mortensen, B., Danielson, B., Harpole, S.W., Alberti, J., Arnillas, C.A., Biederman, L., et al. (2017). Herbivores safeguard plant diversity by reducing variability in dominance. J. Ecol., 38, 42–49.

O’Mara, F.P. (2012). The role of grasslands in food security and climate change. Ann. Bot., 110, 1263–1270.

Olff, H., De Leeuw, J., Bakker, J.P., Platerink, R.J. & Van Wijnen, H.J. (1997). Vegetation Succession and Herbivory in a Salt Marsh: Changes Induced by Sea Level Rise and Silt Deposition Along an Elevational Gradient. J. Ecol., 85, 799–814.

Olff, H. & Ritchie, M.E. (1998). Effects of herbivores on grassland plant diversity. Trends Ecol. Evol., 13, 261–265.

Pétillon, J., Ysnel, F., Canard, A. & Lefeuvre, J.C. (2005). Impact of an invasive plant (Elymus athericus) on the conservation value of tidal salt marshes in western France and implications for management: Responses of spider populations. Biol. Conserv., 126, 103–117.

Reijs, J.W., Daatselaar, J.W.C.H.G., Helming, J.F.M., Jager, J. & Beldman, a C.G. (2013). Grazing dairy cows in North-West Europe. LEI Wageningen UR, Hague.

R Core Team (2019). R: A language and environment for statistical computing. R Foundation for Statistical Computing, Vienna, Austria. URL https://www.R-project.org/.

Ripple, W.J., Newsome, T.M., Wolf, C., Dirzo, R., Everatt, K.T., Galetti, M., et al. (2015). Collapse of the world’s largest herbivores. Sci. Adv., 1, e1400103.

Rupprecht, F., Wanner, A., Stock, M. & Jensen, K. (2015). Succession in salt marshes - large-scale and long-term patterns after abandonment of grazing and drainage. Appl. Veg. Sci., 18, 86–98.

Schrama, M., Heijning, P., Bakker, J.P., Van Wijnen, H.J., Berg, M.P. & Olff, H. (2013a). Herbivore trampling as an alternative pathway for explaining differences in nitrogen mineralization in moist grasslands. Oecologia, 172, 231–243.

Schrama, M., Veen, G.F.C., Bakker, E.S.L., Ruifrok, J.L., Bakker, J.P. & Olff, H. (2013b). An integrated perspective to explain nitrogen mineralization in grazed ecosystems. Perspect. Plant Ecol. Evol. Syst., 15, 32–44.

Tälle, M., Deák, B., Poschlod, P., Valkó, O., Westerberg, L. & Milberg, P. (2016). Grazing vs. mowing: A meta-analysis of biodiversity benefits for grassland management. Agric. Ecosyst. Environ., 222, 200–212.

Terres, J.M., Scacchiafichi, L.N., Wania, A., Ambar, M., Anguiano, E., Buckwell, A., et al. (2015). Farmland abandonment in Europe: Identification of drivers and indicators, and development of a composite indicator of risk. Land use policy, 49, 20–34.

Thornton, P.K. (2010). Livestock production: Recent trends, future prospects. Philos. Trans. R. Soc. B Biol. Sci., 365, 2853–2867.

Tscharntke, T., Clough, Y., Wanger, T.C., Jackson, L., Motzke, I., Perfecto, I., et al. (2012). Global food security, biodiversity conservation and the future of agricultural intensification. Biol. Conserv., 151, 53–59.

Veeneklaas, R.M., Dijkema, K.S., Hecker, N. & Bakker, J.P. (2013). Spatio-temporal dynamics of the invasive plant species Elytrigia atherica on natural salt marshes. Appl. Veg. Sci., 16, 205–216.

Vivian-smith, G. (1997). Microtopographic Heterogeneity and Floristic Diversity in Experimental Wetland Communities. J. Ecol., 85, 71–82.

Wanner, A., Suchrow, S., Kiehl, K., Meyer, W., Pohlmann, N., Stock, M., et al. (2014). Scale matters: Impact of management regime on plant species richness and vegetation type diversity in Wadden Sea salt marshes. Agric. Ecosyst. Environ., 182, 69–79.

Wood S (2017). Generalized Additive Models: An Introduction with R, 2 edition. Chapman and Hall/CRC.

