## Supplementary file for long-term non-trophic effects for "Long-term non-trophic effects of large herbivores on plant diversity are underestimated"

**Supplementary Text**

*Aboveground biomass*

Mowing was executed yearly since 1972. However, only 28 occasions of recordings (till 2016) containing biomass data for all three mowing treatments were available. In addition, vegetation was surveyed in the early season mowing, and the both early and late season mowing for 17 occasions during the 46-year experiment (1972, 1974-1980, 1984-1989, 2003, 2015 and 2017). We usually mowed in late June or early July for the early season mowing, and in late August or early September for the late season mowing. We measured aboveground biomass from the mown plots by first weighing the fresh weight of the cut vegetation. We then took a subsample from the total plant material collected from each mown plot, and determined fresh and dry weight. The fresh: dry ratio was used to calculate the aboveground biomass for mown plots in g dw $m^{-2}$. In August 1982, we measured aboveground biomass from the ungrazed control and the grazing treatment by clipping vegetation of 5 randomly selected 20 cm ×20 cm plots, adjacent to the permanent plots. Biomass from these 5 plots per permanent plot was added up, and multiplied by 5 to estimate the g dw $m^{-2}$ for the ungrazed control and the grazing treatment. In September 2018, before the late season mowing, we measured aboveground biomass for all the treatments. We clipped vegetation of two randomly chosen strips (10 cm × 100 cm) to the ground level (ca. 1 cm) adjacent to the permanent plots, and weighed the biomass to the nearest 0.01 g after drying in the oven (70 °C) to constant weight. The biomass from two strips per permanent plot was added up, and multiplied by 5 to estimate the g dw $m^{-2}$ for all the treatments (data presented in Fig. S2B).

*Relationships between temperature, rainfall, sea level change and plant diversity*

We downloaded the temperature and rainfall data from www.knmi.nl, and sea level change data from https://beeldbank.rws.nl/. We selected years to match those in which vegetation surveys were conducted. To reduce the dependence of sampling in time and space, we first calculated changes in temperature, rainfall and sea level as the values of a given year minus that of the previous year. We calculated change in plant diversity for each permanent plot in a similar way, and we averaged change in plant diversity across all treatments. We calculated the pearson correlation for changes in temperature, rainfall and sea level and plant diversity, respectively.

*Elytrigia atherica and Featuca rubra, and the relationship between dominance and plant diversity*

We explored the percent cover of *E. atherica* and *F. rubra*, which were the most dominant plant species in the ungrazed control and mowing treatment, respectively, 46 years after the start of the experiment. We explored the relationship between dominance and plant diversity, as the current theory suggests reducing dominance is one of the main mechanisms by which large herbivores promote plant diversity (Koerner *et al.* 2018). To reduce the dependence of sampling in time and space, we first calculated changes in plant diversity and dominance for each permanent plot as plant diversity (or dominance) for a given year minus that of the previous year. We averaged the values across four blocks for each treatment, and calculated the pearson correlation for changes in plant diversity and dominance for each treatment.

*Functional groups*

We recorded 30 forb, 13 graminoid, 3 legume and 2 woody species (Table S1). Averaged over all years, large herbivores increased 3.6 forbs and 2.4 graminoids, while non-trophic effects contributed 2 and 1.13 to these increases, respectively (Table S2). In addition, large herbivores promoted forbs and graminoids over time, and the contribution of non-trophic effects increased over time (Fig. S4A, B, C, D; Table S3). Averaged over all years, large herbivores increased the abundance of forbs by 9 %, while they decreased the abundance of graminoids by 7 %, both mainly attributed to the non-trophic effects (Table S2). In addition, large herbivores increased the abundance of forbs over time, which was also mainly attributed to the non-trophic effects (Fig. S5A, B). However, large herbivores decreased the abundance of graminoids over time, which was attributed more to the non-trophic effects before 2000, but more to the trophic effects after 2000 (Fig. S5C, D). Large herbivores increased the abundance of legumes after 2000, which was mainly attributed to the non-trophic effects (Fig. S5E, F).

We recorded 19 rare, 17 frequent, 4 common and 8 abundant species (Table S1). Averaged over all years, large herbivores significantly increased 1.93 rare, 1.2 frequent, 1.57 common, and 1.27 abundant species. Non-trophic effects significantly contributed 1.04 and 1.25 plant species to the increase in rare and common species, respectively (Table S2). In addition, large herbivores promoted rare, frequent, and common species over time. The contribution of non-trophic effects increased in frequent and common species over time (Fig. S6, Table S3).

*Bareground*

As previous studies show that bare ground increases germination and plant diversity (Bakker 1985; Bakker *et al.* 1985), we explored percent cover of bare ground over time. Percent cover of are ground was estimated as 100 - total percent cover of living plants. As we estimated percent cover for each species independently, total cover of living plants can sometimes exceed 100 for the multilayer canopies. In these situations, percent cover of bare ground was defined as 0.

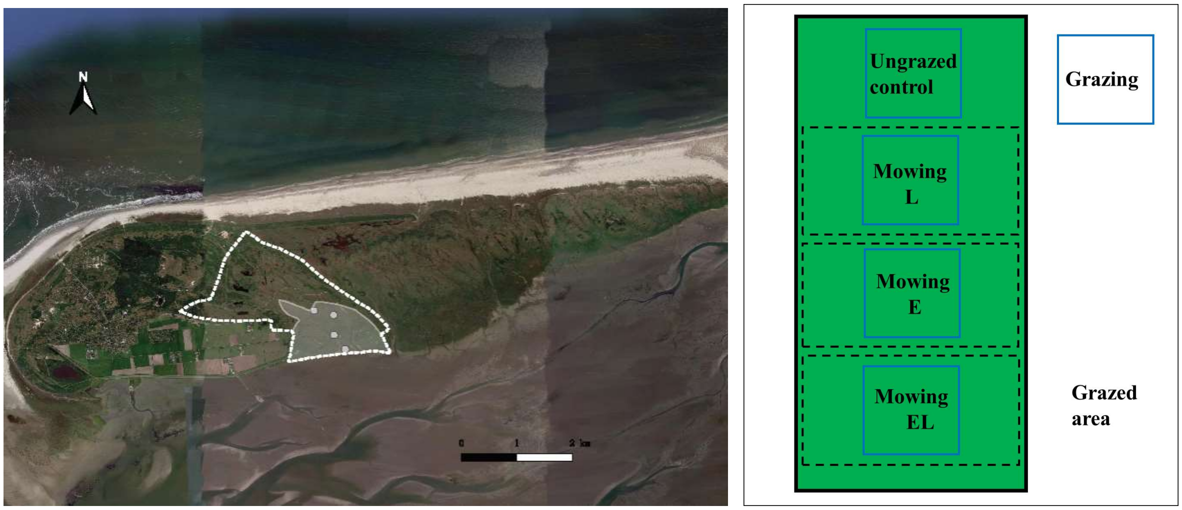


**Fig. S1.**

**Description of study site (left panel) and experimental set-up (right panel).** The drawn grey area is under cattle grazing since 1972, grazing expanded to the dotted white area since 1993. The four white dots represent the four blocks. Treatments within each block are shown in the right panel. Dashed black rectangles were subjected to different mowing treatments, permanent plots (blue rectangles) were established within treatments. In the field, the treatments were randomized within a block. Size of the exclosures and permanent plots were not projected according to their actual measurements. Details can be found in the supplementary text. E: early season mowing, EL: both early and late season mowing, L: late season mowing.

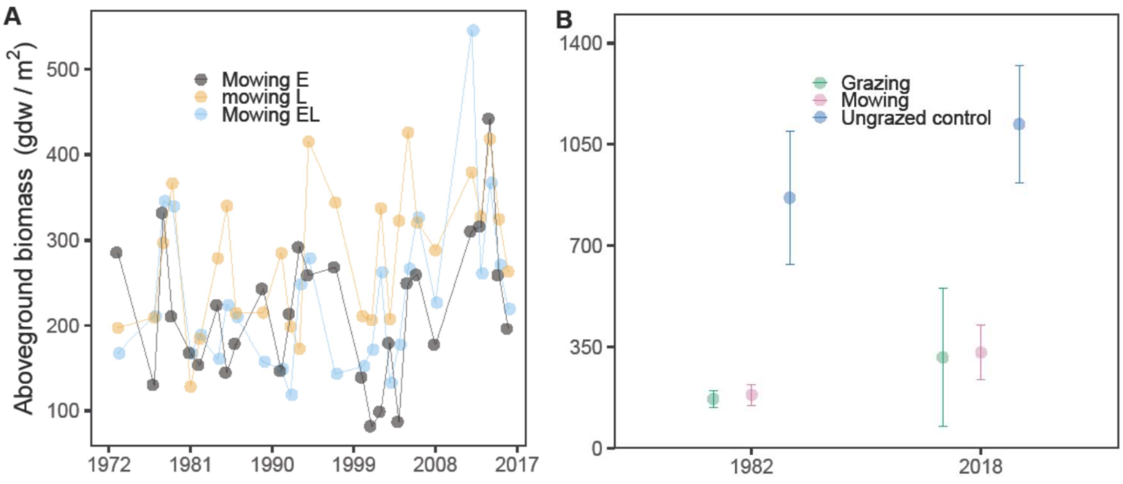


**Fig. S2.**

**Aboveground biomass**. The late season mowing removed the largest amount of aboveground biomass (17 of 28 recordings) over time compared with the early season mowing, and the both early and late season mowing (A). The late season mowing removed a similar amount of aboveground biomass compared with cattle grazing (B). Dots are the means of four blocks. Bars reflect ±1se. E: early season mowing, EL: both early and late season mowing, L: late season mowing.

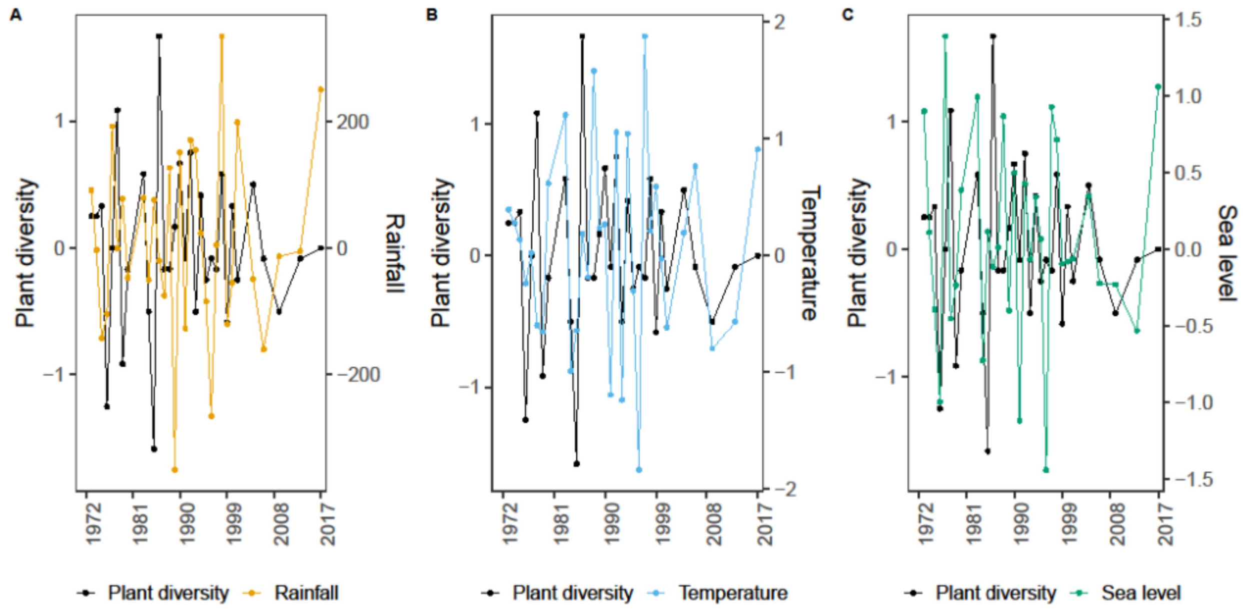


**Fig. S3.**

**Relationships between changes in plant diversity and rainfall, temperature, sea level change.** Change in plant diversity was not correlated with rainfall (Pearson correlation coefficient = 0.56, p = 0.5804), temperature (Pearson correlation coefficient = 1.75, p = 0.0897), and sea level (Pearson correlation coefficient = 1.31, p = 0.1994).

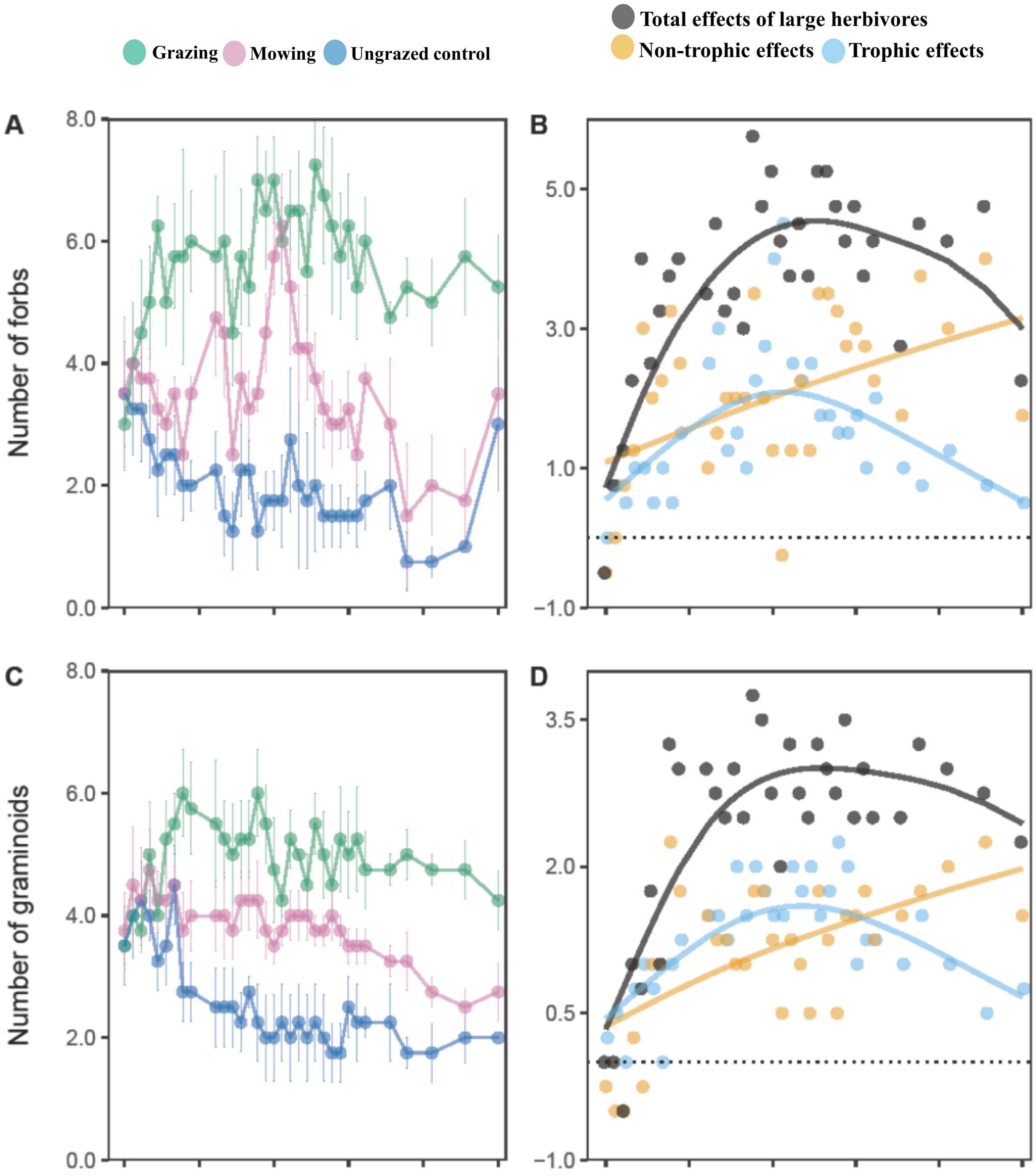
Grazing
Mowing
Ungrazed control
Total effects of large herbivores
Non-trophic effects
Trophic effects
A
B
C
D
Number of forbs
Number of graminoids
8.0
6.0
4.0
2.0
0.0
5.0
3.0
1.0
−1.0
3.5
2.0
0.5
−1.0

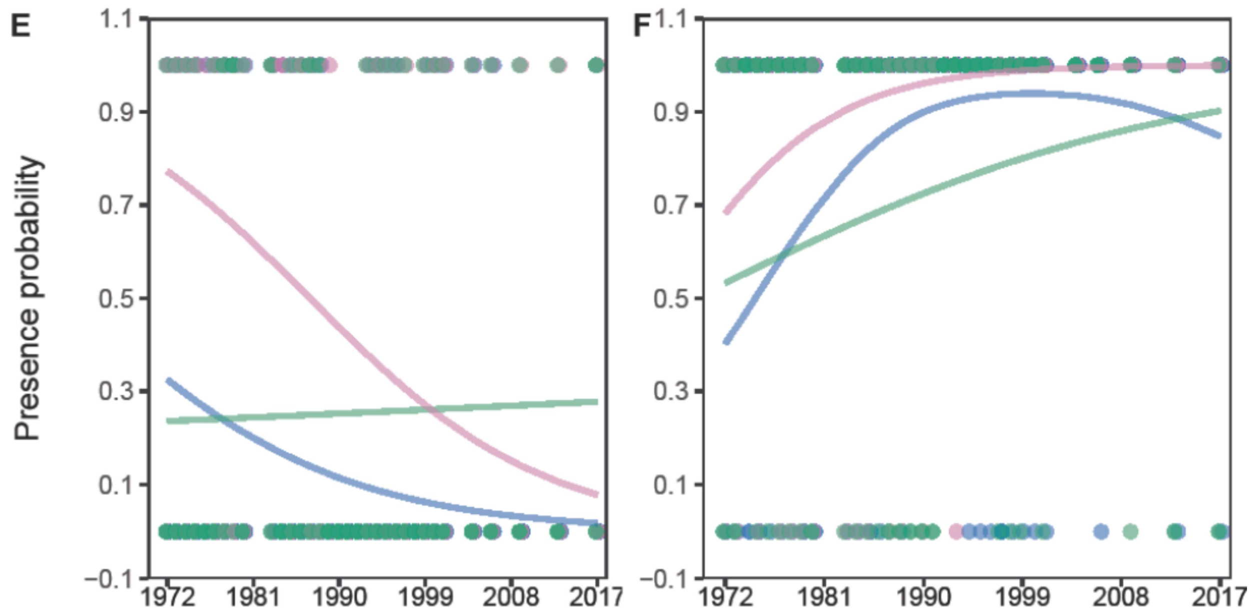


**Fig. S4.**

**Number of forbs (A, B), graminoids (C, D), presence probability of legumes (E) and woody species (F) in the 4-m$^2$ permanent plots.** Large herbivores promoted forbs and graminoids over time, and the contribution of non-trophic effects increased over time (A, B, C, D). Total effects of large herbivores: the grazing treatment minus the ungrazed control; non-trophic effects: the grazing treatment minus the mowing treatment; trophic effects: the mowing treatment minus the ungrazed control. Dots show means of four blocks. Bars reflect ±1se. Lines in B, D, E, F were fitted with gamm models (Table S3).

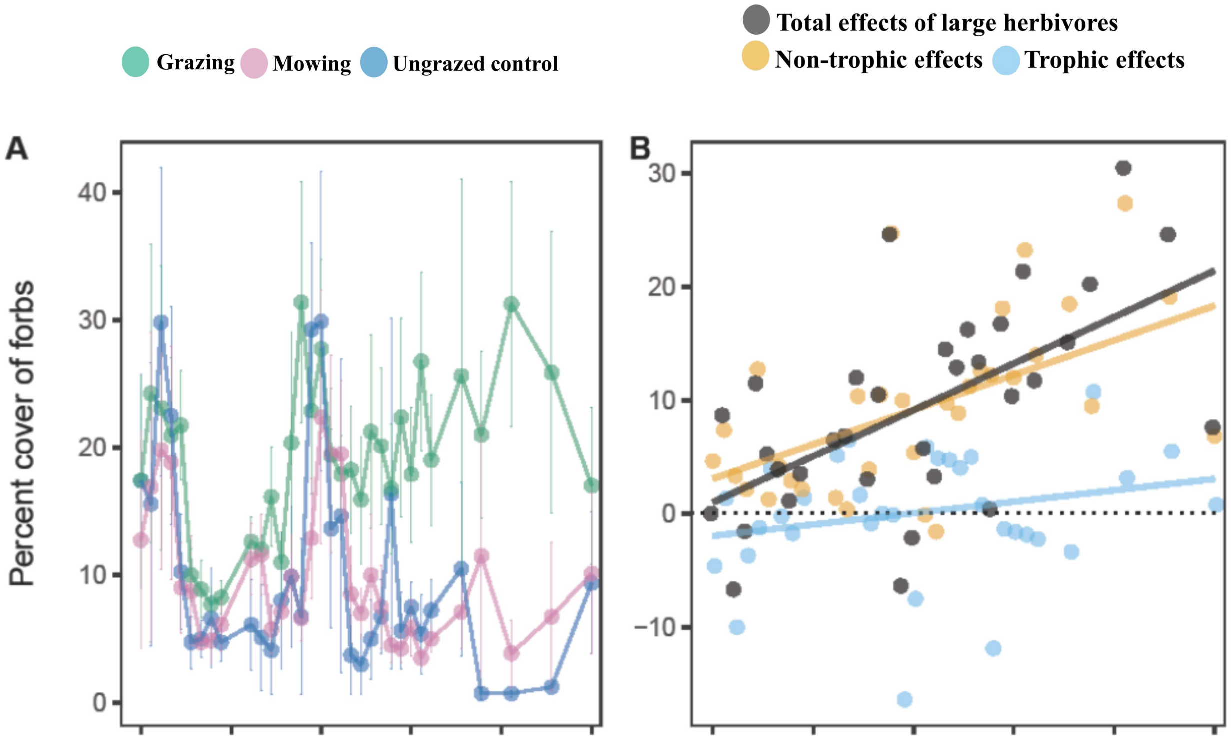
Grazing
Mowing
Ungrazed control
Total effects of large herbivores
Non-trophic effects
Trophic effects
A
Percent cover of forbs
40
30
20
10
0
B
30
20
10
0
−10

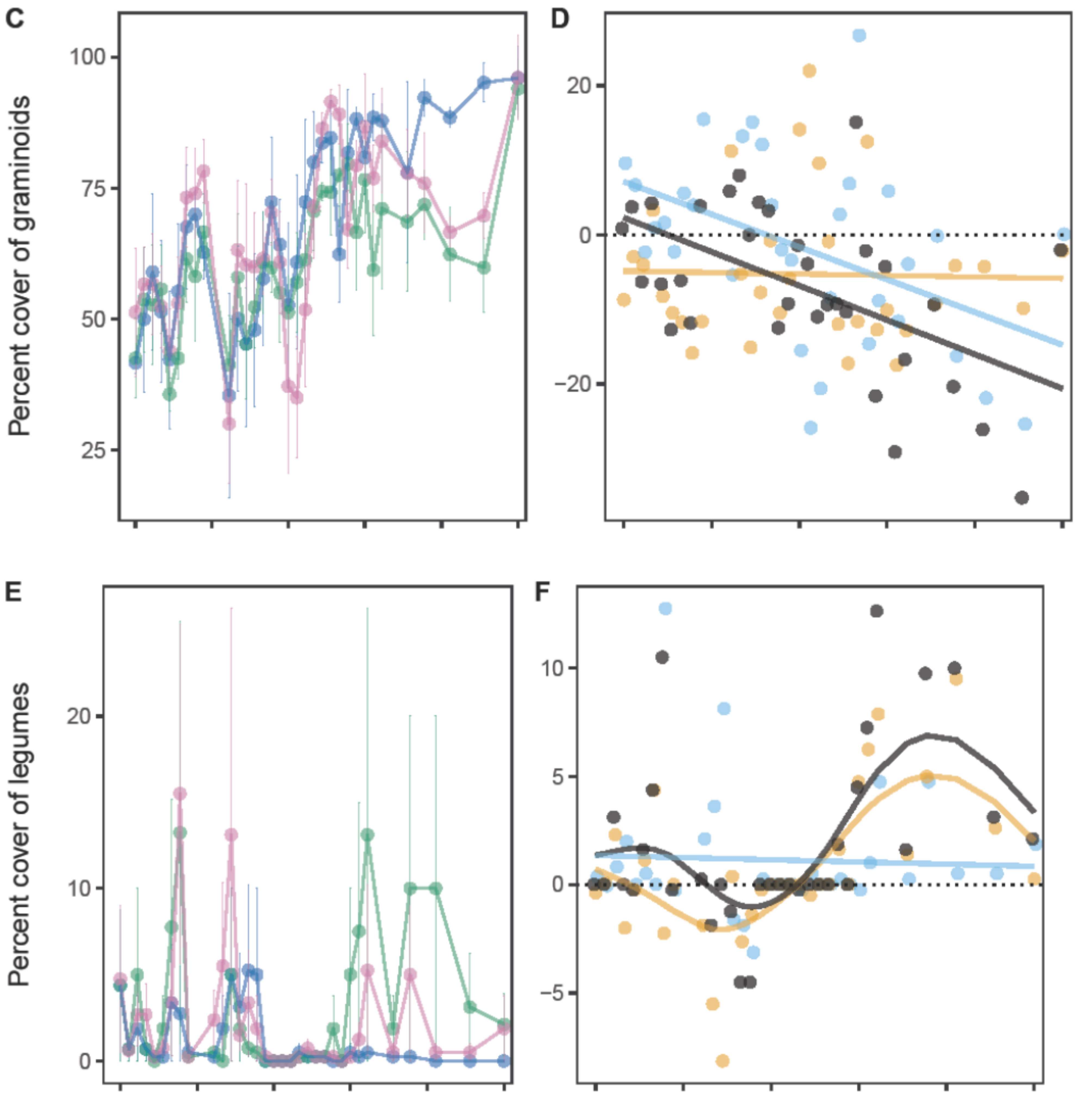
C
D
E
F
Percent cover of graminoids
Percent cover of legumes
100
75
50
25
20
0
−20
20
10
0
10
5
0
−5

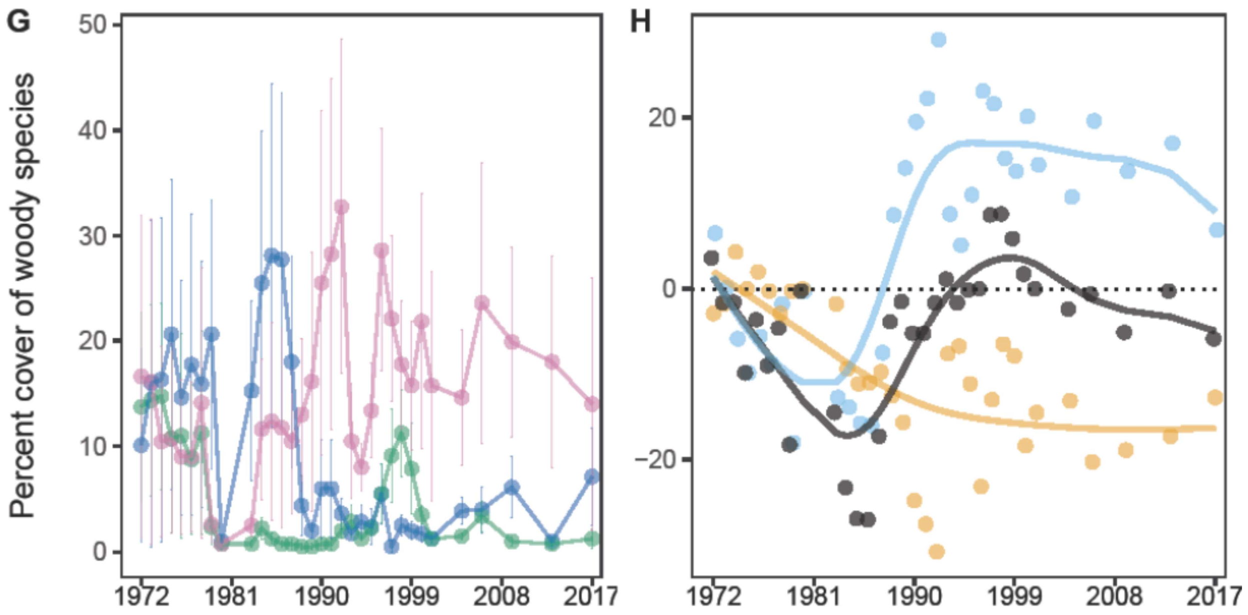


**Fig. S5.**

**Percent cover of forbs (A, B), graminoids (C, D), legumes (E, F) and woody species (G, H).** Large herbivores promoted the abundance of forbs over time, and this was mainly attributed to the non-trophic effects (A, B). Large herbivores decreased the abundance of graminoids over time, which was more attributed the non-trophic effects before 2000, afterwards it was more attributed to the trophic effects (C, D). Total effects of large herbivores: the grazing treatment minus the ungrazed control; non-trophic effects: the grazing treatment minus the mowing treatment; trophic effects: the mowing treatment minus the ungrazed control. Dots show means of four blocks. Bars reflect ±1se. Lines in B, D, F, H are fitted with gamm models (Table S3).

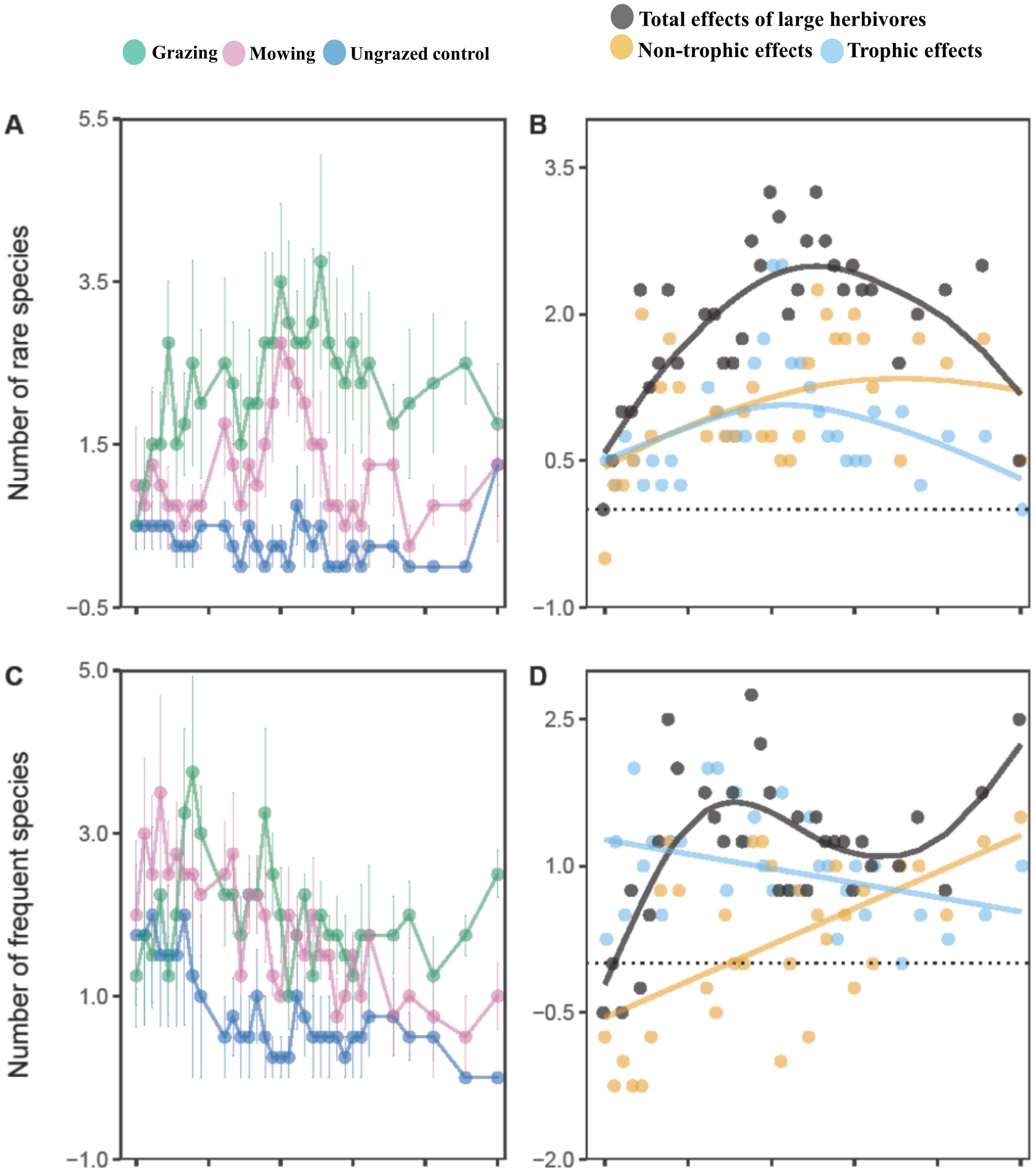
Total effects of large herbivores
Non-trophic effects
Trophic effects
Grazing
Mowing
Ungrazed control
A
B
C
D
Number of rare species
Number of frequent species
5.5
3.5
1.5
−0.5
3.5
2.0
0.5
−1.0
5.0
3.0
1.0
−1.0
2.5
1.0
−0.5
−2.0

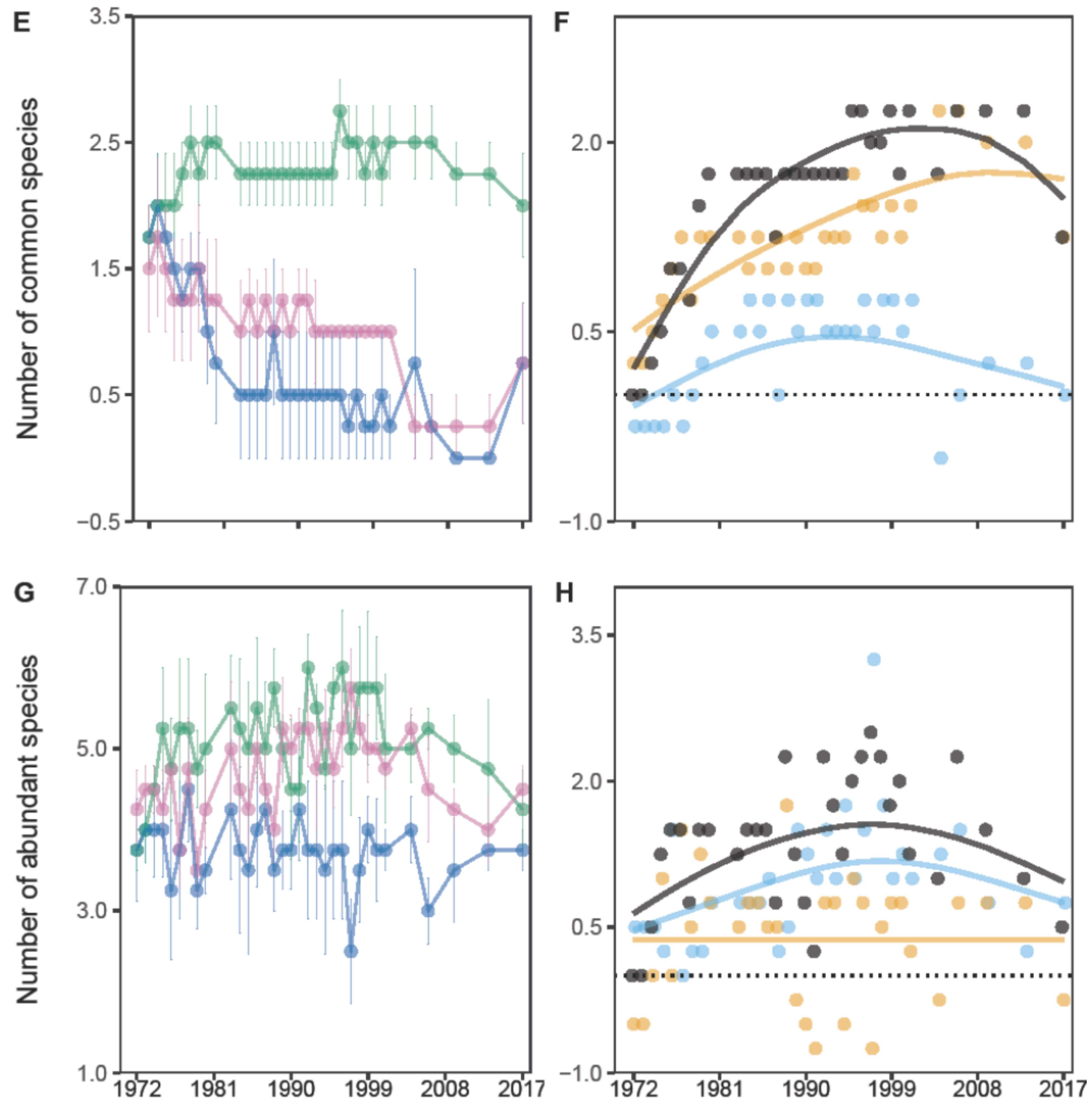


**Fig. S6.**

**Number of rare (A, B), frequent (C, D), common (E, F) and abundant (G, H) species in the 4-m$^2$ permanent plots.** Large herbivores promoted rare, frequent, and common species over time. The contribution of non-trophic effects increased in frequent and common species. Total effects of large herbivores: the grazing treatment minus the ungrazed control; non-trophic effects: the grazing treatment minus the mowing treatment; trophic effects: the mowing treatment minus the ungrazed control. Dots show means of four blocks. Bars reflect 1se. Lines in B, D, F, H were fitted with gamm models (Table S3).

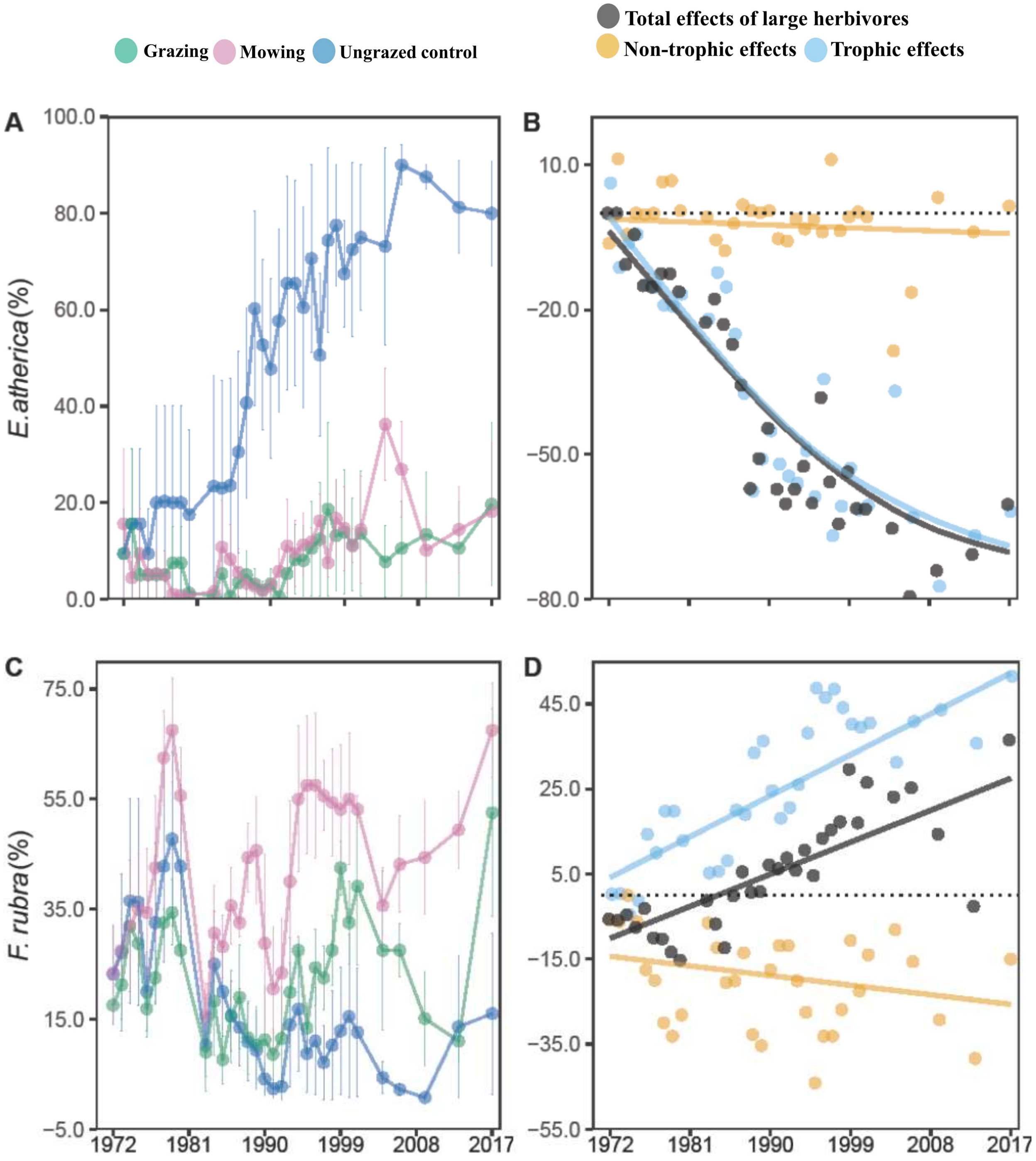

**Fig. S7.**

***Elytrigia atherica* (A, B) and *Festuca rubra* (C, D)**. Large herbivores suppressed the expansion of *E. atherica* over time*,* which was almost entirely attributed to trophic effects. Large herbivores increased the expansion of *F. rubra,* largely attributed to trophic effects. Total effects of large herbivores: the grazing treatment minus the ungrazed control; non-trophic effects: the grazing treatment minus the mowing treatment; trophic effects: the mowing treatment minus the ungrazed control. Dots show means of four blocks. Bars reflect ±1se. Lines in B, D were fitted with gamm models (Table S3).

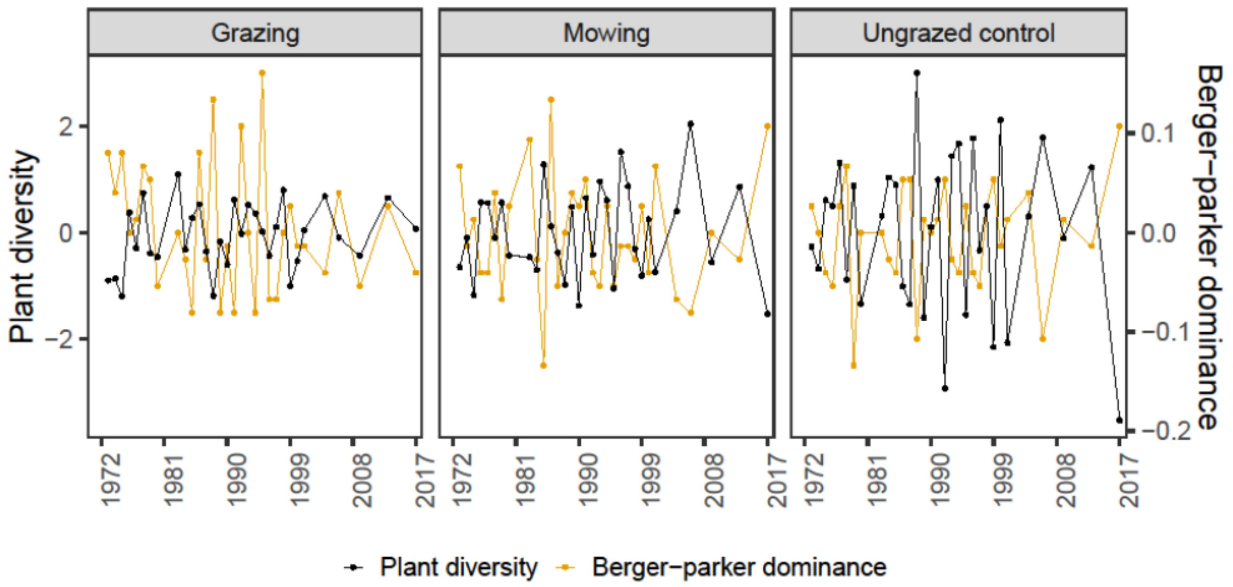


**Fig. S8.**

**Relationship between plant diversity and dominance**. Dominance and plant diversity were negatively correlated, particularly in the ungrazed control treatment (ungrazed control: Pearson correlation coefficient = - 0.71, $p < 0.0001$; grazing: Pearson correlation coefficient = - 0.25, $p = 0.1653$; mowing: Pearson correlation coefficient = - 0.47, $p = 0.0070$).

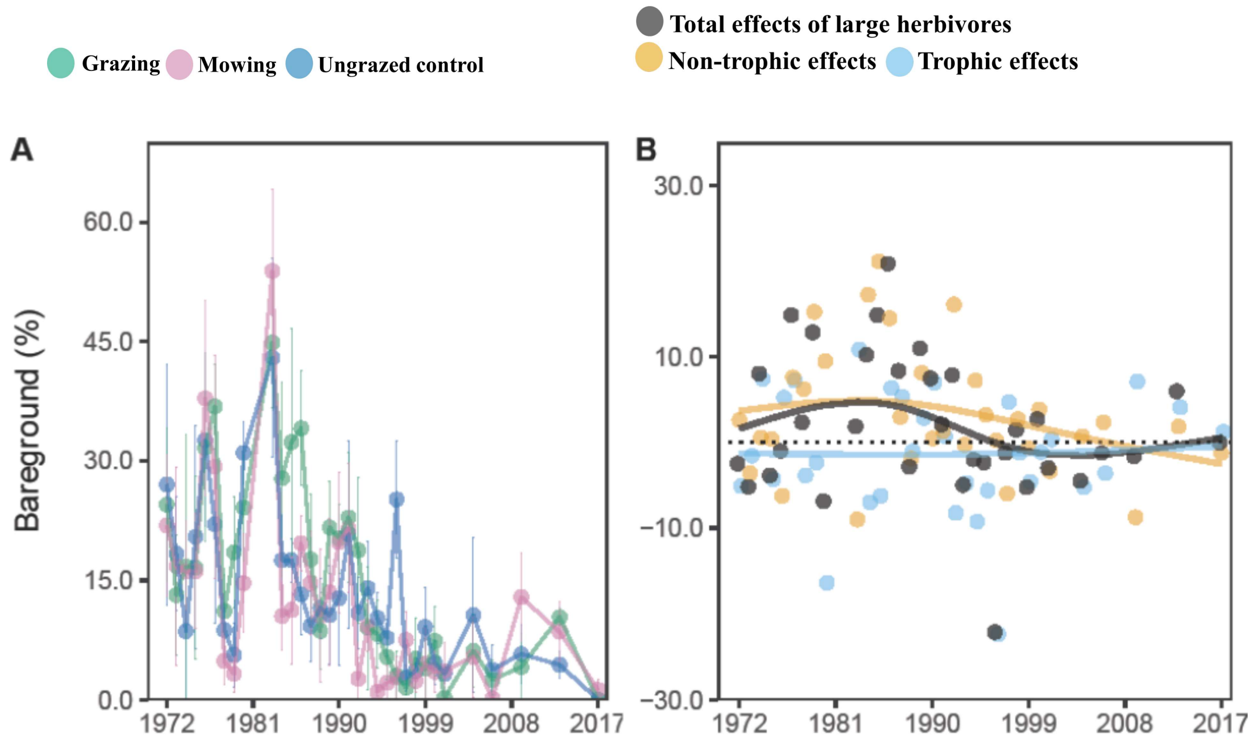


**Fig. S9.**

**Bare ground**. Large herbivores did not significantly impact percent cover of bareground. In the first half of the experiment, large herbivores slightly increased percent cover of bareground, which was primarily attributed to non-trophic effects. Total effects of large herbivores: the grazing treatment minus the ungrazed control; non-trophic effects: the grazing treatment minus the mowing treatment; trophic effects: the mowing treatment minus the ungrazed control. Dots show means of four blocks. Bars reflect ±1se. Lines in B were fitted with gamm models.

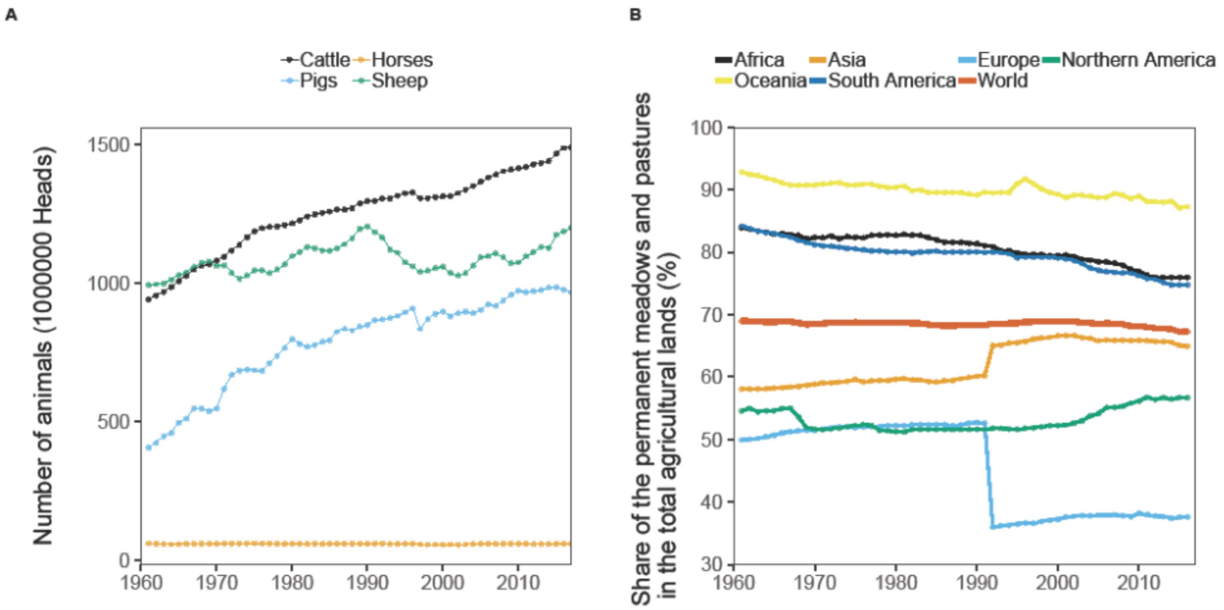


**Fig. S10.**

**Number of livestock (A) and share of permanent meadows and pastures in the total agriculture lands (B).** Numbers of cattle, pigs and sheep increased by 58 %, 138 % and 21 %, respectively, in the last 50 years globally (A). Share of permanent meadows and pastures in the total agriculture lands in the world remained relatively stable. However, this share decreased by more than 15 % after 1992 in Europe, it also decreased in other regions such as Africa, Oceania, and South America (B). Permanent meadows and pastures refer to land used (> 5 years) to grow herbaceous forage crops through cultivation or naturally (wild prairie or grazing land) (FAO; Food and Agriculture Organization). Regions were classified according to the FAO. Date from FAOSTAT (http://www.fao.org/faostat/en/#data).

**Table S1.**

List of species occurring during the 46-year experiment. Names of species, their functional traits and presence/absence in the ungrazed control, grazing and mowing treatment are given. Species in boldface belong to the halophytes, according to (Bakker *et al.* 2002). Rare: percent cover in any permanent plot in any year ≤ 1; frequent: > 1 and ≤ 20; common: > 20 and ≤ 50; abundant: > 50. Life form was classified according to https://wilde-planten.nl. Presence and absence in different treatments (ungrazed control, grazing, mowing), 1: present; 0: absent.

| Species | Abundance | Life form | Presence |
|---|---|---|---|
| *Agrostis stolonifera* | abundant | graminoid | (1, 1, 1) |
| *Armeria maritima* | frequent | forb | (1, 1, 1) |
| *Artemisia maritima* | abundant | woody | (1, 1, 1) |
| *Aster tripolium* | rare | forb | (1, 1, 1) |
| *Atriplex littoralis* | rare | forb | (1, 1, 1) |
| *Atriplex portulacoides* | frequent | woody | (1, 1, 1) |
| ***Atriplex prostrata*** | abundant | forb | (1, 1, 1) |
| *Bromus hordeaceus* ssp. *hordeaceus* | frequent | graminoid | (1, 0, 1) |
| *Bupleurum tenuissimum* | rare | forb | (0, 1, 1) |
| *Carex distans* | frequent | graminoid | (1, 1, 1) |
| *Centaurium pulchellum* | rare | forb | (0, 1, 1) |
| *Cerastium fontanum* ssp. *vulgare* | rare | forb | (1, 1, 1) |
| *Cirsium arvense* | frequent | forb | (0, 0, 1) |
| *Cochlearia danica* | frequent | forb | (0, 1, 1) |

| | | | |
|---|---|---|---|
| *Cochlearia officinalis* ssp. *anglica* | frequent | forb | (1, 1, 1) |
| *Elytrigia atherica* | abundant | graminoid | (1, 1, 1) |
| *Elytrigia repens* | rare | graminoid | (0, 1, 0) |
| *Festuca rubra* | abundant | graminoid | (1, 1, 1) |
| ***Glaux maritima*** | common | forb | (1, 1, 1) |
| *Juncus gerardii* | abundant | graminoid | (1, 1, 1) |
| *Juncus maritimus* | common | graminoid | (1, 1, 1) |
| *Leontodon autumnalis* | rare | forb | (0, 0, 1) |
| *Limonium vulgare* | rare | forb | (1, 1, 1) |
| *Lotus corniculatus* | frequent | legume | (1, 0, 1) |
| *Odontites vernus* ssp. *serotinus* | abundant | forb | (1, 1, 1) |
| *Parapholis strigosa* | frequent | graminoid | (0, 1, 1) |
| *Plantago coronopus* | rare | forb | (0, 1, 1) |
| *Plantago lanceolata* | rare | forb | (0, 1, 1) |
| ***Plantago maritima*** | common | forb | (1, 1, 1) |
| *Poa annua* | frequent | graminoid | (1, 1, 0) |
| *Poa pratensis* | frequent | graminoid | (1, 1, 1) |
| *Potentilla anserina* | common | forb | (1, 1, 1) |
| ***Puccinellia maritima*** | abundant | graminoid | (1, 1, 1) |
| *Sagina maritima* | rare | forb | (0, 1, 1) |
| *Sagina nodosa* | rare | forb | (0, 1, 1) |
| *Sagina procumbens* | rare | forb | (1, 0, 0) |
| ***Salicornia* spp.** | rare | forb | (1, 1, 1) |

| | | | |
|---|---|---|---|
| *Sonchus arvensis* | rare | forb | (0, 0, 1) |
| ***Spergularia media*** | rare | forb | (1, 1, 1) |
| ***Spergularia salina*** | frequent | forb | (0, 1, 1) |
| *Stellaria graminea* | frequent | forb | (0, 1, 1) |
| *Stellaria media* | rare | forb | (0, 1, 0) |
| ***Suaeda maritima*** | rare | forb | (1, 1, 1) |
| *Taraxacum* spp. | frequent | forb | (0, 0, 1) |
| *Trifolium pratense* | frequent | legume | (0, 1, 0) |
| *Trifolium repens* | frequent | legume | (1, 1, 1) |
| *Triglochin maritima* | frequent | graminoid | (1, 1, 1) |
| *Tripleurospermum maritimum* | rare | forb | (0, 0, 1) |

**Table S2.**

Parameters estimated from the gamm models. For each variable, results from two models were included, one for the total effects of large herbivores, the other for the trophic and non-trophic effects. For the results of the trophic and non-trophic effects, the significance of non-trophic effects was assessed by the first p value, the significance between the trophic and non-trophic effects was assessed by the second p value.

| Variables | Parameters | Estimated (Mean ±1se) | t.value | *p* |
|---|---|---|---|---|
| Plant diversity | Large herbivores | 5.91 ± 1.234 | 4.79 | < 0.0001 |
| | Non-trophic effects | 2.81 ± 1.002 | 2.8 | 0.0054 |
| | Trophic effects | 2.99 ± 1.419 | 0.12 | 0.902 |
| Cumulative species gain | Large herbivores | 0.23 ± 0.054 | 4.21 | < 0.0001 |
| | Non-trophic effects | 0.13 ± 0.098 | 1.33 | 0.1854 |
| | Trophic effects | 0.09 ± 0.138 | -0.27 | 0.7895 |
| Cumulative species loss | Large herbivores | -0.31 ± 0.073 | -4.32 | < 0.0001 |
| | Non-trophic effects | -0.17 ± 0.134 | -1.25 | 0.2137 |
| | Trophic effects | -0.16 ± 0.190 | 0.06 | 0.9513 |
| Forbs | Large herbivores | 3.60 ± 0.705 | 5.11 | < 0.0001 |
| | Non-trophic effects | 2.00 ± 0.615 | 3.25 | 0.0013 |
| | Trophic effects | 1.55 ± 0.871 | -0.52 | 0.6064 |
| Graminoids | Large herbivores | 2.39 ± 0.594 | 4.03 | 0.0001 |
| | Non-trophic effects | 1.13 ± 0.342 | 3.29 | 0.0011 |
| | Trophic effects | 1.23 ± 0.485 | 0.22 | 0.8292 |
| Legumes | Ungrazed control | -2.07 ± 1.333 | -1.55 | 0.1223 |
| | Grazing | -1.08 ± 0.618 | 1.59 | 0.1133 |
| | Mowing | -0.28 ± 0.615 | 2.9 | 0.004 |
| Woody | Ungrazed control | 1.68 ± 1.051 | 1.6 | 0.111 |

| | | | | |
|---|---|---|---|---|
| | Grazing | 0.98 ± 1.020 | -0.68 | 0.4954 |
| | Mowing | 3.24 ± 1.192 | 1.31 | 0.1912 |
| Percent cover of forbs | Large herbivores | 9.31 ± 2.335 | 3.99 | 0.0001 |
| | Non-trophic effects | 9.29 ± 3.202 | 2.9 | 0.004 |
| | Trophic effect | 0.07 ± 4.528 | -2.04 | 0.0427 |
| Percent cover of graminoids | Large herbivores | -7.02 ± 2.928 | -2.4 | 0.0179 |
| | Non-trophic effects | -5.24 ± 3.105 | -1.69 | 0.0926 |
| | Trophic effect | -1.78 ± 4.391 | 0.79 | 0.4318 |
| Percent cover of legumes | Large herbivores | 1.81 ± 1.744 | 1.04 | 0.3001 |
| | Non-trophic effects | 0.68 ± 0.919 | 0.74 | 0.4613 |
| | Trophic effect | 1.14 ± 1.300 | 0.35 | 0.7246 |
| Percent cover of woody species | Large herbivores | -4.89 ± 2.832 | -1.73 | 0.0865 |
| | Non-trophic effects | -10.51 ± 4.002 | -2.63 | 0.0091 |
| | Trophic effect | 5.62 ± 5.659 | 2.85 | 0.0047 |
| Dominance | Large herbivores | -0.20 ± 0.054 | -3.64 | 0.0004 |
| | Non-trophic effects | -0.07 ± 0.056 | -1.18 | 0.2372 |
| | Trophic effects | -0.13 ± 0.079 | -0.79 | 0.4325 |
| Rare species | Large herbivores | 1.93 ± 0.587 | 3.28 | 0.0013 |
| | Non-trophic effects | 1.04 ± 0.445 | 2.35 | 0.0197 |
| | Trophic effects | 0.85 ± 0.629 | -0.31 | 0.7565 |
| Frequent species | Large herbivores | 1.20 ± 0.222 | 5.4 | < 0.0001 |
| | Non-trophic effects | 0.21 ± 0.344 | 0.6 | 0.5478 |
| | Trophic effects | 0.97 ± 0.487 | 1.57 | 0.1183 |
| Common species | Large herbivores | 1.57 ± 0.212 | 7.42 | < 0.0001 |
| | Non-trophic effects | 1.25 ± 0.177 | 7.08 | < 0.0001 |
| | Trophic effects | 0.29 ± 0.250 | -3.85 | 0.0002 |
| Abundant | Large herbivores | 1.27 ± 0.549 | 2.32 | 0.0219 |

| | | | | |
|---|---|---|---|---|
| species | Non-trophic effects | 0.37 ± 0.299 | 1.23 | 0.2184 |
| | Trophic effects | 0.93 ± 0.424 | 1.32 | 0.1882 |
| *E.atherica* | Large herbivores | -38.73 ± 8.198 | -4.72 | < 0.0001 |
| | Non-trophic effects | -2.44 ± 6.022 | -0.41 | 0.6857 |
| | Trophic effects | -37.36 ± 8.563 | -4.08 | 0.0001 |
| *F.rubra* | Large herbivores | 5.23 ± 10.598 | 0.49 | 0.6226 |
| | Non-trophic effects | -18.95 ± 7.411 | -2.56 | 0.0111 |
| | Trophic effects | 23.77 ± 10.481 | 4.08 | 0.0001 |
| Halophytes | Large herbivores | 2.93 ± 0.598 | 4.9 | < 0.0001 |
| | Non-trophic effects | 2.41 ± 0.367 | 6.57 | < 0.0001 |
| | Trophic effects | 0.51 ± 0.519 | -3.66 | 0.0003 |
| Percent cover of halophytes | Large herbivores | 13.83 ± 4.422 | 3.13 | 0.0022 |
| | Non-trophic effects | 16.23 ± 3.679 | 4.41 | < 0.0001 |
| | Trophic effect | -2.18 ± 5.191 | -3.54 | 0.0005 |

**Table S3.**

Temporal trends (smooths) estimated from the gamm models. For each variable, results from two models were included, one for the total effects of large herbivores, the other for the trophic and non-trophic effects.

| Variables | Smoothers | edf | F | *p* | $R^2$ |
|---|---|---|---|---|---|
| Plant diversity | Large herbivores | 2.76 | 3.21 | 0.02 | 0.157 |
| | Non-trophic effects | 1.68 | 2.52 | 0.0547 | 0.072 |
| | Trophic effects | 1.93 | 1.09 | 0.3549 | 0.072 |
| Cumulative species gain | Large herbivores | 2.52 | 2.61 | 0.1181 | 0.044 |
| | Non-trophic effects | 1 | 1.28 | 0.2581 | 0.019 |
| | Trophic effects | 1.6 | 4.74 | 0.04 | 0.019 |
| Cumulative species loss | Large herbivores | 2.11 | 7 | 0.0011 | 0.232 |
| | Non-trophic effects | 1 | 10.82 | 0.0011 | 0.07 |
| | Trophic effects | 2.61 | 3.8 | 0.0221 | 0.07 |
| Forbs | Large herbivores | 2.91 | 3.84 | 0.0089 | 0.144 |
| | Non-trophic effects | 1.14 | 3.7 | 0.0407 | 0.076 |
| | Trophic effects | 2.29 | 2.27 | 0.1511 | 0.076 |
| Graminoids | Large herbivores | 2.83 | 5.54 | 0.0043 | 0.191 |
| | Non-trophic effects | 1.35 | 4.97 | 0.0123 | 0.102 |
| | Trophic effects | 2.36 | 2.75 | 0.0841 | 0.102 |
| Legumes | Ungrazed control | 1 | 4.25 | 0.0401 | 0.012 |
| | Grazing | 1 | 0.02 | 0.8829 | 0.012 |
| | Mowing | 1 | 6.85 | 0.0093 | 0.012 |
| Woody | Ungrazed control | 2.05 | 4.91 | 0.0065 | 0.093 |
| | Grazing | 1 | 3 | 0.0841 | 0.093 |
| | Mowing | 1 | 6.67 | 0.0103 | 0.093 |
| Percent cover of forbs | Large herbivores | 1 | 21.84 | < 0.0001 | 0.146 |
| | Non-trophic effects | 1 | 9.84 | 0.0019 | 0.156 |
| | Trophic effect | 1 | 1.06 | 0.3043 | 0.156 |
| Percent cover of graminoids | Large herbivores | 1 | 5.61 | 0.0193 | 0.052 |
| | Non-trophic effects | 1 | 0.01 | 0.9138 | 0.022 |
| | Trophic effect | 1 | 5.23 | 0.023 | 0.022 |
| Percent cover of legumes | Large herbivores | 3.93 | 4.1 | 0.0083 | 0.084 |
| | Non-trophic effects | 3.78 | 5.8 | 0.0006 | 0.061 |
| | Trophic effect | 1 | 0.06 | 0.8001 | 0.061 |
| Percent cover of woody | Large herbivores | 5.33 | 7.96 | < 0.0001 | 0.216 |

| | | | | | |
|---|---|---|---|---|---|
| species | Non-trophic effects | 2.28 | 8.73 | 0.0001 | 0.321 |
| | Trophic effect | 5.4 | 11.18 | < 0.0001 | 0.321 |
| Dominance | Large herbivores | 2.15 | 7.39 | 0.0007 | 0.113 |
| | Non-trophic effects | 1 | 6.1 | 0.0141 | 0.095 |
| | Trophic effects | 2.2 | 2.31 | 0.1117 | 0.095 |
| Rare species | Large herbivores | 2.79 | 4.55 | 0.0072 | 0.111 |
| | Non-trophic effects | 1.73 | 1.07 | 0.2202 | 0.038 |
| | Trophic effects | 1.97 | 1.31 | 0.2906 | 0.038 |
| Frequent species | Large herbivores | 3.62 | 4.26 | 0.0069 | 0.17 |
| | Non-trophic effects | 1 | 13.29 | 0.0003 | 0.121 |
| | Trophic effects | 1 | 2.07 | 0.1514 | 0.121 |
| Common species | Large herbivores | 2.94 | 5.67 | 0.0012 | 0.306 |
| | Non-trophic effects | 1.99 | 5.24 | 0.0078 | 0.325 |
| | Trophic effects | 2.04 | 1.14 | 0.2925 | 0.325 |
| Abundant species | Large herbivores | 1.8 | 0.69 | 0.4228 | 0.028 |
| | Non-trophic effects | 1 | 0 | 0.9951 | 0.057 |
| | Trophic effects | 1.86 | 0.84 | 0.3459 | 0.057 |
| *E.atherica* | Large herbivores | 1.75 | 5.04 | 0.0061 | 0.353 |
| | Non-trophic effects | 1 | 0.06 | 0.8 | 0.499 |
| | Trophic effects | 2.01 | 18.38 | < 0.0001 | 0.499 |
| *F.rubra* | Large herbivores | 1 | 6.32 | 0.0131 | 0.144 |
| | Non-trophic effects | 1 | 0.96 | 0.3288 | 0.501 |
| | Trophic effects | 1 | 17.34 | < 0.0001 | 0.501 |
| Halophytes | Large herbivores | 4.09 | 8.81 | < 0.0001 | 0.243 |
| | Non-trophic effects | 5.82 | 4.16 | 0.0007 | 0.408 |
| | Trophic effects | 2.8 | 4.21 | 0.017 | 0.408 |
| Percent cover of halophytes | Large herbivores | 2.67 | 5.61 | 0.0015 | 0.205 |
| | Non-trophic effects | 3.07 | 5.96 | 0.0006 | 0.395 |
| | Trophic effect | 1 | 0.81 | 0.3689 | 0.395 |
